# Cell cycle follows “pause and play” mechanism in environment stress recovery in diverse plant species

**DOI:** 10.1101/2025.04.02.646900

**Authors:** Olivia S. Hazelwood, Kamryn A. Diehl, Vicky Hollenbeck, Dustin Herb, Joseph P. Gallagher, M. Arif Ashraf

**Affiliations:** Department of Botany, University of British Columbia, Vancouver, BC, Canada; Forage Seed and Cereal Research Unit, United States Department of Agriculture, Corvallis, OR, USA

**Keywords:** Abiotic stress, Stress recovery, Cell cycle, AlphaFold, Plant development, Root growth

## Abstract

Changes to organismal growth induced by environmental stress are orchestrated at the cellular level. These periods of stress may be followed by recovery periods, when plants have the opportunity to return to normal growth conditions. However, the cell cycle mechanisms underlying recovery are poorly understood. We tested the cell cycle regulation in roots during control, stress, and recovery period for salt, osmotic, cold, and heat stresses using *Arabidopsis thaliana*, *Brachypodium distachyon* and *Lolium multiflorum*. During the salt and cold stress conditions, the cell cycle pauses at gap phase and is released from gap phase during stress recovery, which depends on CDKA;1 and ICK1. Cold stress and recovery, which specifically relies on cell division, follows a conserved “pause and play” mechanism of cell cycle.

---

Environmental stresses, such as salinity, drought, cold, and heat, alter plant growth and development (*1*, *2*). Plant growth is a combination of cell division, cell elongation, and cell differentiation (*3*, *4*). The reduction in growth during environmental stress response alters either the cell division, cell elongation, or both (*4*, *5*). How distinct environmental stresses target either cell division or elongation is still being unraveled. Furthermore, stress adaptation relies not only on survival during the stress period, but also on a plant’s capacity to recover after stress conditions. Plant growth during stress and stress recovery is orchestrated at the cellular level and manifests at the tissue and organ level. Cell cycle regulators control the cellular circuits important for both cell division and elongation and switch between these core functions and integrating environmental cues (*6*). A comparative genetics and cell biology approach for a wide range of environmental stresses and recovery could illustrate a fundamental mechanism important for plant adaptation, and, more broadly, an adaptive stress response for organisms. This understanding of stress response and recovery will allow us to connect the cellular regulators to the plant phenotypes they control.

## Distinctive environmental stress and stress recovery response in plants

We investigated the root growth response of Arabidopsis (*Arabidopsis thaliana*) during environmental stresses and recovery periods. Firstly, we used the 4-day-old Arabidopsis root to measure both primary root length and cell length during the stress and recovery period (Fig. 1, A to F, and fig. S1). We observed ∼40%, ∼50%, 80% primary root growth inhibition for salt (48h), osmotic (mannitol) stress (48h), and cold (4°C) stress (24h), respectively, and 10% acceleration of primary root growth during heat (29°C) stress (24h) (Fig. 1, A to F, and fig. S1). It is not surprising that salt, osmotic, and cold stress inhibit the primary root growth, as numerous studies have demonstrated a similar phenomenon (*4*, *7–11*). At the same time, high temperature induced primary root growth acceleration has also been observed for Arabidopsis and other plant root systems (*12–15*).

**Figure 1:**
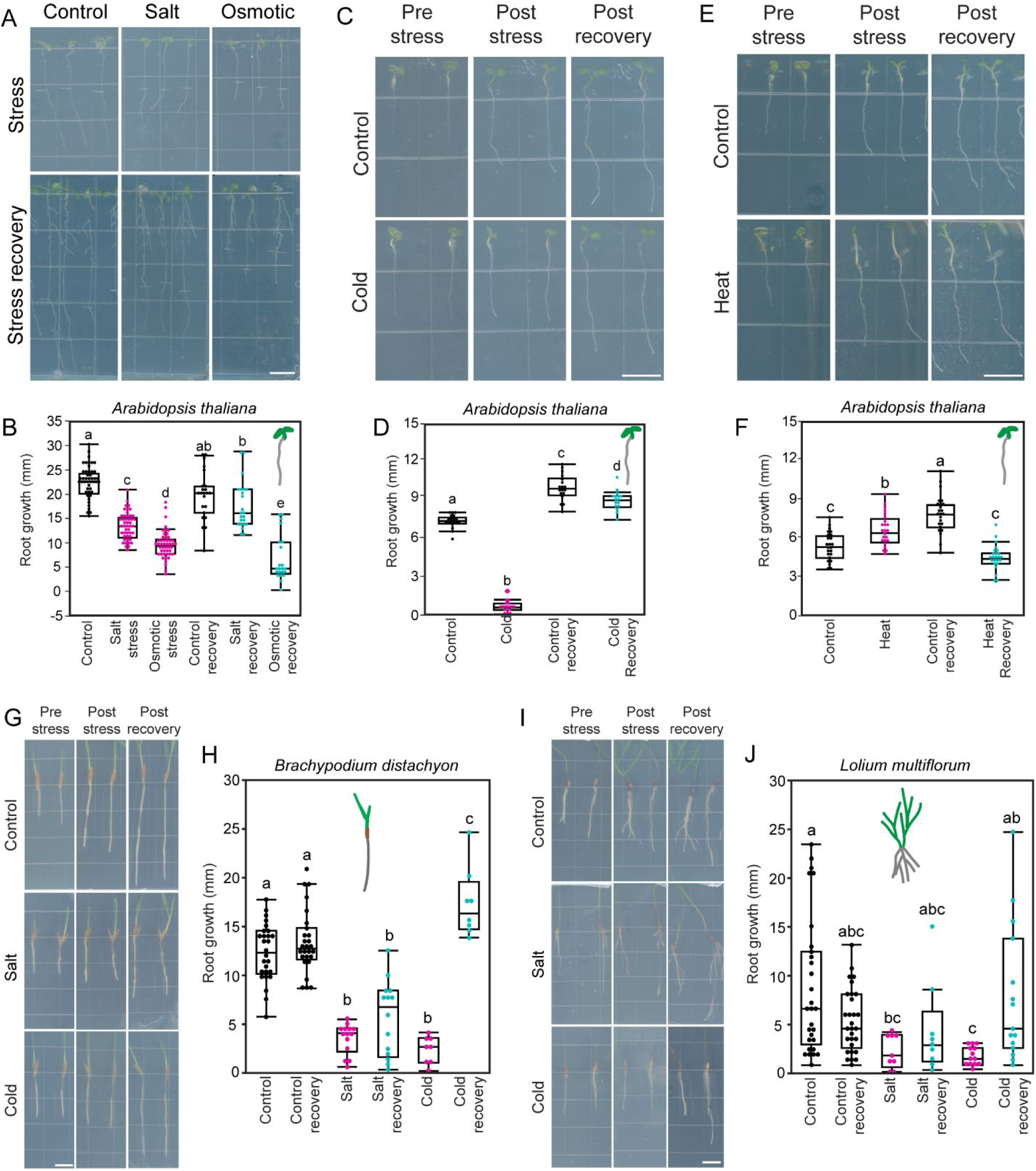
Environmental stress and stress recovery in Arabidopsis, Brachypodium, and annual ryegrass. (A) Root growth response of Arabidopsis in presence of salt and mannitol stress (upper panel) and recovery (lower panel). (B) Quantification of root growth from A. (C) Root growth response of Arabidopsis during pre-and post-cold stress, and post-cold stress recovery conditions. (D) Quantification of root growth from C. (E) Root growth response of Arabidopsis during pre-and post-heat stress, and post-heat stress recovery conditions. (F) Quantification of root growth from E. (G) Examples of root growth response of Brachypodium during salt and cold stress, and consequent recovery period. (H) Quantification of Brachypodium root growth in salt and cold stress and recovery experiments. (I) Examples of root growth response of annual ryegrass during salt and cold stress, and consequent recovery period. (J) Quantification of annual ryegrass root growth in salt and cold stress and recovery experiments.In panels A, C, E, G, and I, scale bar = 10 mm. Statistical test is performed based on Tukey’s Honest test (B, D, F, H, and J). Groups labeled with the same letter are not statistically different from each other (alpha = 0.05). Boxplots show median values (center line), 25th to 75th interquartile range (box) and 1.5*interquartile range (whiskers) (B, D, F, H, and J).

In contrast to stress conditions, the outcome during the recovery period is surprising. Given an equal amount of time to recover following the stress condition, Arabidopsis root growth recovers from salt and cold stress (Fig. 1, A to D, and fig. S1) yet fail to recover from osmotic and heat stresses (Fig. 1, A,B,E,F, and fig. S1). Within 48h of transfer from the salt to control plates, Arabidopsis root starts growing similar to control recovery (Fig. 1, A and B). Similarly, cold-treated Arabidopsis roots require only 24h to recover 90% compared to the control recovery (Fig. 1, C and D). Given the importance of abiotic stress recovery in plants, we hypothesized that this recovery potential may be conserved beyond Arabidopsis. We tested the salt and cold stress recovery in two grasses – Brachypodium (*Brachypodium distachyon*) and annual ryegrass (*Lolium multiflorum*). Overall, root growth was more variable within these two species. Despite this higher variability, root growth rates still recover in Brachypodium and annual ryegrass during salt and cold stress recovery, although only the increase in root growth in cold stress recovery was significant (Fig.1, G to J, and fig. S1). In Brachypodium, the recovery from cold stress (∼4 fold) is greater than salt stress recovery (∼2 fold) (Fig.1, G to H), while in annual ryegrass, a similar salt (∼1.5 fold) and cold (∼1.5 fold) stress recovery pattern was observed (Fig.1, I to J). Because of differences in root architecture across these three species, a conserved cellular mechanism seems highly likely to control these changes in growth under stress and recovery.

### Cell cycle mechanism during salt and cold stress recovery

The visible root growth during stress and recovery period happens through a combination of cell division and elongation (*3–5*). Cell division relies on a tightly regulated cell cycle, consisting of G1 (gap phase 1), S (DNA synthesis phase), G2 (gap phase 2), and M (mitosis). Molecular markers (CYCB1;1-GUS/GFP, Cytrap, PlaCCI, PCNA1-sGFP) in plants have been used for both visualization and quantification of cell cycle phases (*16–20*). In Arabidopsis, we utilized cell cycle markers Cytrap (indicates S/G2 and G2/M phases) and PCNA1-sGFP (highlights gap phase, early and late S) to quantify the cell cycle status and measured epidermal cell length to investigate the cell elongation. Both salt and osmotic stresses reduce the meristematic cell division activity and cell elongation (Fig. 2, A to F, and fig. S2). This data confirms the previous observation of reduced CYCB1;1-GUS activity during salt stress in Arabidopsis root (*21*). Interestingly, during the recovery period, both cell division and elongation, and consequently the root growth, recover for salt stress recovery, but not for osmotic stress recovery (Fig. 1, A to B, Fig. 2, A to F, and fig. S2). Previous studies on Arabidopsis and maize leaf during osmotic stress highlighted the importance of cell division activity as a key factor for growth revival during the rehydration stage (*22*, *23*).

**Figure 2:**
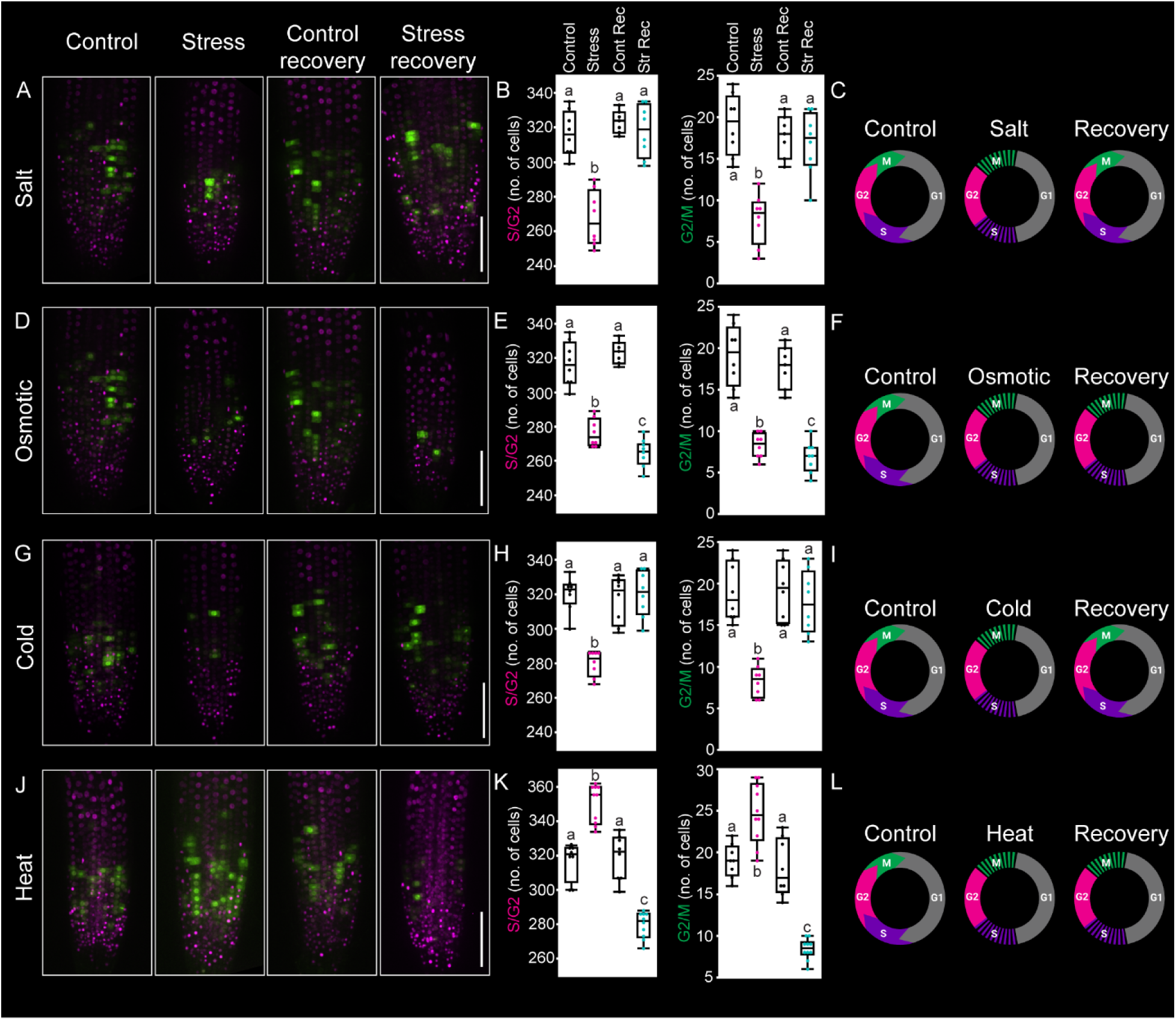
Response of cell cycle marker during environmental stress and stress recovery in Arabidopsis. Representative images of cell cycle marker Cytrap, quantification of S/G2 and G2/M phase specific cells, and cell cycle regulation model during salt (A-C), osmotic (D-F), cold (G-I), and heat (J-L) stress and recovery periods. Magenta and green indicate S/G2 and G2/M phase, respectively, specific cells. At least 8-10 roots were observed for each condition and the entire Z plane of the root meristem was considered to quantify the cells. Scale bar = 100 µm (A, D, G, and J). Statistical test is performed based on Tukey’s Honest test (B, E, H, and K). Groups labeled with the same letter are not statistically different from each other (alpha = 0.05). Boxplots show median values (center line), 25th to 75th interquartile range (box) and 1.5*interquartile range (whiskers) (B, E, H, and K).

Previously, we demonstrated that decreased root growth in cold stress is due to a decrease in G2/M phase cell cycle progression, and not due to a decrease in cell elongation(*4*). At the same time, more cells were observed in gap phase, visualized using PCNA1-sGFP, during cold stress (*4*). Here, we have observed that the cold stress recovery period is sufficient for root growth recovery, as well as to resume the cell cycle (Fig. 2, G to I, and fig. S3). Compared to other stress conditions, heat stressed plants demonstrate a faster root growth and an accelerated cell cycle during heat stress; however, they fail to recover like control conditions during the recovery period (Fig. 2, J to L, and fig. S3). Altogether, our data suggest that salt and cold stress recovery for root growth is tightly correlated with the cell cycle regulation (Fig. 2, A to L, and fig. S4). We further tested this concept using PCNA1-sGFP during salt stress and recovery period. Reminiscent of cold stress, more cells were observed in gap phase during salt stress, yet they resumed normal cell cycle patterns during the salt stress recovery (fig. S5). In Brachypodium, a similar pattern of reduced mitotic activity is observed due to salt stress (*24*). This quantitative cell biology approach during the stress and recovery period led us to conclude that the cell cycle follows a “pause and play” mechanism during the salt and cold stress recovery. Between salt and cold stress, the former alters both cell division and cell elongation and the latter only affect cell division, not elongation (fig. S4). Because only one aspect of growth is affected by cold stress, we hypothesized that cell cycle-mediated cold stress recovery will follow a more conserved mechanism across diverse plant species compared to salt stress.

### Salt and cold stress recovery require intact G1 to S and G2 to M transitions

To explore which genes might be regulating the observed changes in the cell cycle, we performed transcriptomics of Arabidopsis whole root tissue during stress (salt, osmotic, cold, and heat) and recovery conditions (fig. S6, table S1). According to principal components analysis, stress sample gene expression is typically distinct from control and recovery sample gene expression (fig. S7). As expected, a low number of genes are differentially expressed between the stress control and recovery control, while hundreds to thousands of genes are differentially expressed between stress and recovery conditions (fig. S6, table S2). Interestingly, a cluster of core Arabidopsis cell cycle genes is differentially expressed during salt and cold stress and recovery (Fig. 3A, and fig. S8). In terms of differential expression of core cell cycle genes, cold stress has a distinct pattern compared to salt stress (Fig. 3A). This supports our data and hypothesis that cold stress impacts only cell division during stress and recovery, but salt stress may incorporate both cell division and cell elongation.

**Figure 3:**
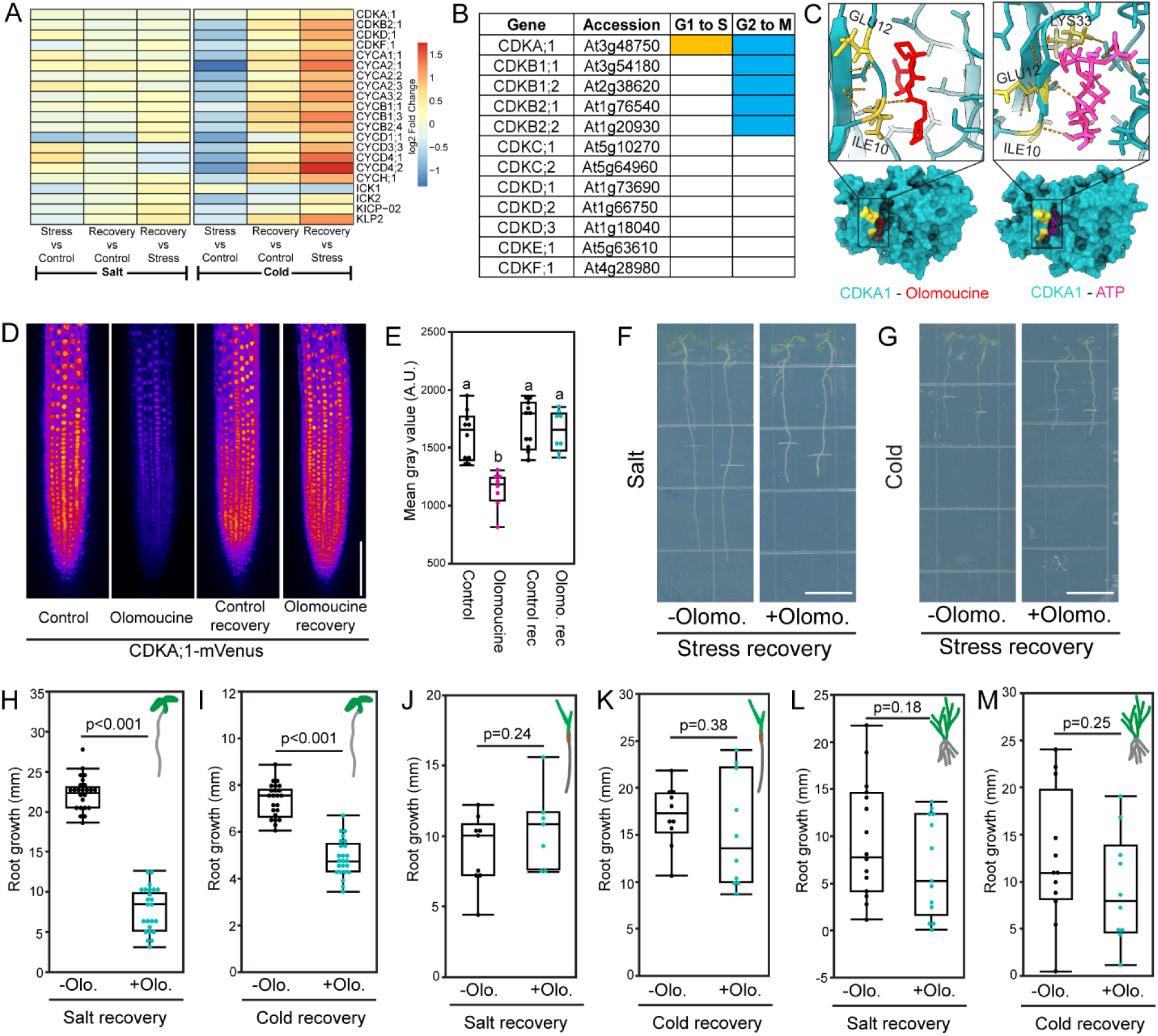
Intact G1 to S and G2 to M transitions are required for salt and cold stress recovery. (A) Relative expression of a sub-set of core cell cycle genes during the salt and cold stress, and subsequent recovery periods. (B) List of Arabidopsis CDKs involved in G1 to S (yellow) and G2 to M (cyan) transitions. (C) *In silico* interaction assay of olomoucine (left, red) and ATP (right, magenta) with CDKA1 (cyan). Interacting amino acids were highlighted in yellow. (D) CDKA;1-mVenus signal in presence of olomoucine and the recovery period. (E) Quantification of mean gray value from D. Salt (F) and cold (G) stress recovery in presence of olomoucine in Arabidopsis. Quantification of salt (H) and cold (I) stress recovery in presence of olomoucine from F and G, respectively. Salt (J) and cold (K) stress recovery in presence of olomoucine in Brachypodium. Salt (L) and cold (M) stress recovery in presence of olomoucine in annual ryegrass. Scale bar = 100 µm (D) and 10 mm (F and G). (E) Statistical test is performed based on Tukey’s Honest test. Groups labeled with the same letter are not statistically different from each other (alpha = 0.05). Boxplots show median values (center line), 25th to 75th interquartile range (box) and 1.5*interquartile range (whiskers). Student’s t-test was performed for H, I, J, K, L, and M.

The core cell cycle genes are primarily divided into three major categories–cyclin dependent kinases (CDKs), cyclins, and inhibitors (ICKs/KRPs) (*6*, *25*). CDKs, cyclins, and ICKs act in the cell cycle in a phase-dependent manner. Relying on the existing literature to classify the role of Arabidopsis core cell cycle proteins during G1 to S and G2 to M transitions (*6*), we identified one CDK (CDKA;1) and two cyclins (CYCD1;1 and CYCD4;1) involved in both G1 to S and G2 to M transitions (Fig. 3B, and fig. S9).

Cyclins express transiently during specific cell cycle phases and degrade after that period. In contrast, CDKs remain stable during the cell cycle phase. As a result, we evaluated the role of CDKA;1 to block both G1 to S and G2 to M transitions.

Unfortunately, *cdka;1* homozygous mutant causes defect during the male gametophyte development (*26*).

To circumvent this issue, we took a chemical genetic approach and used olomoucine, which inhibits CDKs in non-plant systems (fig. S10) (*27–29*). Before using the olomoucine to block the cell cycle, we tried to understand the specificity of the olomoucine by studying its interaction with CDKA;1 *in silico*. The AlphaFold3 predicted structure of CDKA;1 (pLDDT = 90) was subjected to molecular dynamics simulations in the presence of either ATP or olomoucine to assess whether CDKA;1’s ATP-binding site exhibits nonspecific affinity for other adenine-based ligands like olomoucine (*30*– *33*). Olomoucine binds to the same cleft as ATP in CDKA;1 (Fig. 3C, fig. S11, and movies S1, S2) (*34*, *35*). Additionally, molecular docking simulations predicted binding affinities (Kd) of 4.3μM and 1.3μM with ATP and olomoucine, respectively. This indicates that olomoucine binds more strongly to CDKA;1’s ATP-binding site and may be able to out-compete ATP at lower concentrations. Since ATP binding with CDKA;1 happens in a reversible manner, we hypothesized that CDKA;1 inhibition caused by olomoucine may also act in a reversible manner. We tested this possibility by treating CDKA;1-mVenus/*cdka;1* transgenic line with olomoucine. Interestingly, the CDKA;1-mVenus signal decreases dramatically after the olomoucine treatment and the signal re-appears after a transfer to a control plate during the recovery period (Fig. 3, D and E).

Altogether, these findings demonstrate that olomoucine binds specifically and reversibly to CDKA;1, validating its use as a targeted and reversible controllable tool for inhibiting CDKA;1 activity *in planta*.

We performed the salt and cold recovery experiment in the presence of olomoucine using Arabidopsis, Brachypodium, and annual ryegrass (Fig. 3, F to M). We created a dose response curve for olomoucine and identified a critical concentration (15 µg/mL) to inhibit ∼50% root growth in Arabidopsis (fig. S9) and used this concentration for Arabidopsis, Brachypodium and annual ryegrass experiment. In Arabidopsis, plants treated with olomoucine had their growth reduced by ∼50% and ∼40% in salt and cold stress recovery, respectively (Fig. 3, H to I).

In Brachypodium and annual ryegrass, we see a similar trend of growth reduction caused by olomoucine in the cold stress recovery phase, with the mean root growth rate lower in olomoucine treated plants (Fig. 3, K and M). This data suggests that cold stress recovery requires intact G1 to S and G2 to M transitions of cell cycle similar to Arabidopsis root. In the meantime, the direction of the salt stress recovery pattern is not shared in Brachypodium and annual ryegrass (Fig. 3, J and L). This may suggest that cell elongation factors play a larger role in salt stress and recovery. Cell cycle regulators are highly conserved across diverse plant species compared to the cell elongation regulation. Additionally, root growth in grass systems always demonstrates higher variability and consequently, the drug treatment contributes to this variability (Fig. 1, H and J, Fig. 3, J to M) (*36–38*). Although olomoucine demonstrates a species-specific effect during salt and cold stress recovery, these data suggest that intact G1 to S and G2 to M transitions are required for cell cycle-specific cold stress recovery across plant species.

### CDKA1-dependent cell cycle regulation is required during stress recovery

Olomoucine-mediated CDKA;1 inhibition ensures a break in the G1 to S and G2 to M transitions. But, it is hard to assess whether olomoucine targets any other cell cycle regulators other than CDKA;1. To clarify this, we tried to find an *in vivo* cell cycle inhibitor of CDKA;1. We relied on the previously published root single cell RNA sequencing data and protein interaction data of CDKA;1 (*6*, *39*, *40*). ICK1 is one of the cell cycle inhibitors identified as CDKA;1 interactor (*41*). Spatial expression of *CDKA;1* (required for G1 to S and G2 to M transitions), *CYCB1;1* (G2 to M transition marker gene), and *ICK1* demonstrated that both *CDKA;1* and *CYCB1;1* expression is specifically high in the root apical meristem and reduces in the transitions and elongation zone (Fig. 4, A and B). In contrast, ICK1 expression is lower in the root apical meristem region and higher in the transition and elongation zone (Fig. 4A). These data clearly suggest a spatial regulation of *CDKA;1* and *ICK1* expression during the root development. We performed a real time PCR using Arabidopsis roots to test the gene expression of *CYCB1;1*, *CDKA1*, and *ICK1* during salt and cold stress, and stress recovery conditions (fig. S12). We found that *CYCB1;1* and *ICK1* gene expressions are altered during the stress and recovery period (fig. S12). Compared to *CYCB1;1* and *ICK1* gene expression, the gene expression pattern for *CDKA1* is not straightforward (fig. S12). More importantly, the entire root was used for the RNA isolation and cDNA preparation to perform the qPCR experiment, where the spatial expression data is masked. Combining the existing single cell RNA sequencing data, bulk RNA sequencing data, and qPCR results, we suggest a stress and recovery specific altered gene expression pattern of core cell cycle genes. Additionally, core cell cycle genes, such as *CDKA1* may function at translational level. We hypothesized that overexpression of *ICK1* will help to inhibit CDKA;1 and consequently block G1 to S and G2 to M more specifically (Fig. 4C) (*42*). We tested this hypothesis by using *ICK1* overexpression during salt and cold stress recovery. We found that *ICK1* overexpression fails to recover during salt and cold stress recovery compared to wild-type (Fig. 4, D to G). Altogether, our observation suggests that CDKA;1-dependent cell cycle regulation to maintain proper G1 to S and G2 to M transitions is required for the salt and cold stress recovery.

**Figure 4:**
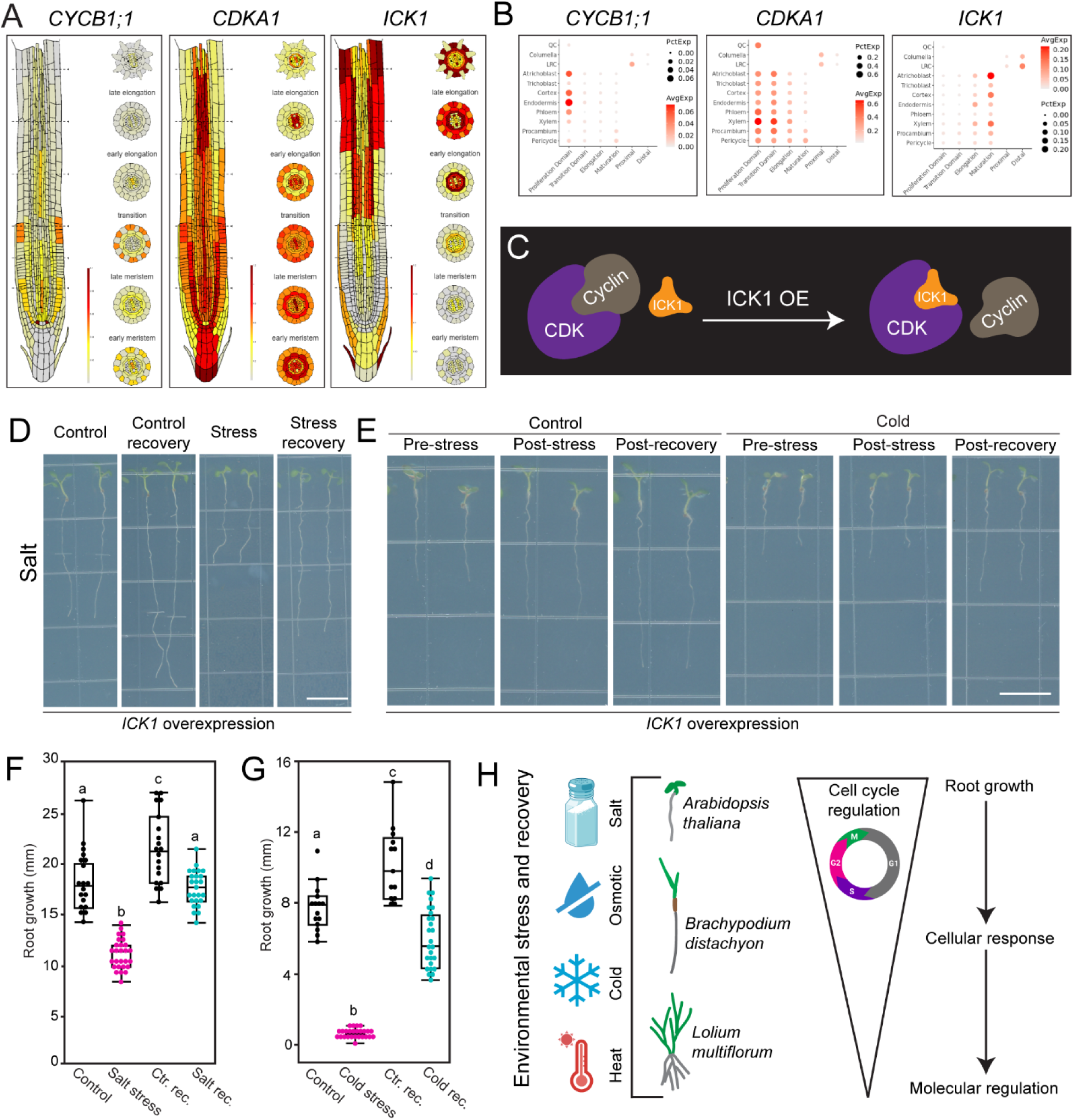
CDKA1-dependent cell cycle transitions are required for salt and cold stress recovery. (A and B) Single cell sequencing data for demonstrating the spatial expression pattern of *CYCB1;1* (left), *CDKA1* (middle), and *ICK1* (right). The upper panel highlights a color based gradient for longitudinal expression pattern and the lower panel highlights the expression pattern in a dot plot manner. (C) Schematic model of ICK1 overexpression based CDK inhibition. Salt (E) and cold (E) recovery response of ICK1 overexpression line in Arabidopsis. Quantification of root growth response of ICK1 overexpression line during salt (F) and cold (G) stress and recovery from E and E, respectively. (H) Graphical model highlights the summarized version of the study, where Arabidopsis, Brachypodium, and annual ryegrass were tested for multiple stress, and stress recovery focusing on cell cycle regulation at organ or root growth, cellular, and molecular scale. Scale bar = 10 mm (D and E). Statistical test is performed based on Tukey’s Honest test (F and G). Groups labeled with the same letter are not statistically different from each other (alpha = 0.05). Boxplots show median values (center line), 25th to 75th interquartile range (box) and 1.5*interquartile range (whiskers) (F and G).

Our study revealed a cell cycle circuit required for environmental stress and recovery mechanisms conserved across Arabidopsis, Brachypodium, and annual ryegrass, species separated by 160 million years of evolution (*43*). This circuit relies on the “pause and play” mechanism of the cell cycle during stress recovery that manifests as root growth (Fig. 4H). Stress recovery is an important aspect of stress resilience that has not been widely considered in past studies, but the growth and performance of plants relies on post-stress recovery (*23*). Therefore this cell cycle regulation is important for both stress and subsequent recovery. We tested the discovery using a wide range of approaches – root growth response, cell biology approach through cell cycle markers, comparative transcriptomics, molecular biology, and chemical genetics (Fig. 4H). This can set a benchmark for studying stress and stress recovery in other plant systems. Understanding root growth response is vital for breeding and engineering crops with tolerance to environmental stresses (*44–47*). Many studies on environmental stresses focus on species-specific stress tolerance. Here, we have found a pathway that is conserved across multiple plant species, and thus is likely conserved in other crop species. This mechanism provides a genetic framework for understanding the network of genes conferring salt and cold tolerance to flowering plants, and potentially other eukaryotic species.

## Supplementary materials

### Materials and methods

#### Plant materials

*Arabidopsis thaliana* accession Columbia (Col-0), *B. distachyon* Bd21-3, and *Lolium mutliflorum* ‘Gulf’ were used as wild types for experiments. The following lines were previously described: Cytrap (*17*), *pAtPCNA1::AtPCNA1-sGFP* (*19*), CDKA;1-mVenus/*cdka;1* (*48*), *35S::GFP-ICK1* (*42*).

#### Growth conditions

For Arabidopsis, seeds were surface sterilized with 1mL 70% ethanol for 10 minutes, washed twice with 1mL autoclaved MQ water inside the laminar flow hood, and placed on the surface of ½ Murashige and Skoog (MS) media (BioWorld; Cat.# 30630058) containing 1% sucrose (BioWorld; Cat.# 41900152) and 1% agar (BioWorld; Cat.# 40100072) in square petri dishes (Simport; Cat.# 26-275). Plate openings were sealed with micropore tapes (Amazon; 3M A-1530-0 Micropore Surgical Tape, ½ Inch), covered with aluminum foil, and kept at 4°C for 2 days. After 2 days, plates are placed vertically under constant light conditions at 22°C.

For *B. distachyon*, seeds were surface sterilized with 0.6% NaOCl, 0.1% Tween-20 for 7 minutes shaking at 200 rpm, washed three times with autoclaved E-pure water inside a laminar flow hood, and left to soak at 4°C for 7 days. Imbibed seeds were placed on the surface of ½ MS media containing 1% sucrose and 1% agar. Plates were sealed with parafilm and oriented vertically under constant light at 22°C.

For *L. multiflorum*, seeds were dehusked in 50% sulfuric acid for 15 minutes, shaking at 200 rpm. Dehusked seeds were rinsed with 500 ml autoclaved E-pure water in a sterile Gooch crucible in the laminar flow hood. Then seeds were sterilized with 3.75% NaOCl, 0.1% Tween-20 for 20 minutes shaking at 200 rpm, rinsed with 500 ml autoclaved E-pure water in a sterile Gooch crucible in the laminar flow hood, and placed on the surface of ½ MS media containing 1% sucrose and 1% agar. Plates were sealed with parafilm and oriented vertically under constant light at 22°C.

#### Stress and stress recovery conditions

Stress and recovery conditions and timeline is summarized in Suppl Fig 1. For the Arabidopsis salt and osmotic stresses, 100mM NaCl and 300mM mannitol-containing ½ MS plates were used, respectively, to incubate seedlings for 2 days. Cold and heat stresses were performed using 4°C and 29°C chambers, respectively, for 1 day period. During the salt and osmotic stress recovery, seedlings were transferred to control media for 2 days. For cold and heat recovery, seedlings were returned to control growth temperature for 1 day period. Standard Arabidopsis growth conditions were maintained in a growth shelf with constant light conditions.

Two days after germination, Brachypodium and annual ryegrass seedlings were incubated in stress conditions for 48 hrs. Seedlings in control conditions were incubated on ½ MS media plates at 22°C under constant light, seedlings under salt stress conditions were incubated on 200 mM NaCl ½ MS media plates under control temperature and light conditions, and seedlings under cold stress conditions were incubated at 4°C in control media and light conditions. After stress conditions, seedlings were moved to new ½ MS media plates at 22°C under constant light for 48 hrs for stress recovery. All experiments took place in Percival or Conviron growth chambers.

#### Phenotypic image analysis

Images of Arabidopsis, Brachypodium, and annual ryegrass root growth were obtained using the Epson Perfection V600 Photo scanner (Ver. 3.9.3.1) at 300 dpi in.TIFF format. Root length was measured using open source ImageJ/Fiji software (*49*). Plant growth was recorded before stress, after stress, and after stress recovery.

#### Drug treatments

Olomoucine (Cayman Chemical, Cat.# 10010240) was dissolved in DMSO and incorporated into ½ MS media at various concentrations for dose response assay.

For olomoucine recovery inhibition experiments, seedlings and stress conditions were the same as described for stress and recovery experiments above. For the stress recovery period, seedlings were either placed on control ½ MS media or ½ MS media with 15 µg/ml olomoucine. Images and measurements were taken as described above.

#### Microscopic imaging

Epidermal cell length was observed using a 20x objective lens attached with a Leica DM 2000 LED compound microscope setup and the images were taken by Leica MC170 HD camera attached with the Leica DM 2000 LED.

Epidermal cell length was also observed with a 20x objective lens attached with AmScope T490 Series Simul-Focal Biological Trinocular Compound Microscope and images were captured using a 18MP USB 3.0 C-mount Camera and AmScope software.

The cell cycle regulation during control, stress, and stress recovery was measured through cell cycle marker, Cytrap, which indicates S/G2 and G2/M states of cell cycle using *pHTR2::CDT1a (C3)-RFP* and *pCYCB1::CYCB1-GFP*, respectively(*15*). The G1, S, and G2 phase of the cell cycle was visualized using *pAtPCNA1::AtPCNA1-sGFP* (*17*) during control, salt stress, and salt recovery conditions. Fluorescent images of live Arabidopsis root meristem were imaged using a Nikon Ti-E-PFS inverted spinning-disk confocal microscope equipped with x20 (Cytrap) and x40 silicone oil immersion (*pAtPCNA1::AtPCNA1-sGFP*) objectives. The system is outfitted with a Yokogawa CSU-X1 spinning disk unit, a self-contained 4-line laser module (excitation at 405, 488, 561, and 640 nm), and Andor iXon 897 EMCCD camera. 488nm and 561nm excitation were used for GFP and RFP, respectively. Fluorescent images were acquired using the Nikon NIS-Elements software and processed using ImageJ software (imagej.nih.gov/ij/). Cells in both the longitudinal (meristematic region) and radial (the entire Z axis) dimensions of root were counted for both Cytrap and *pAtPCNA1::AtPCNA1-sGFP* roots.

CDKA;1-mVenus images were acquired using an Olympus IXplore SpinSR system (Evident, Tokyo, Japan), equipped with an inverted microscope (IX83; Evident, Japan), a CSUW1-Sora spinning disk confocal unit (Yokogawa, Japan) and a Hamamatsu ORCA Fusion BT camera. Fluorescence images were obtained with a 20x objective (UPLXAPO 20X/0.8NA) and fluorescent signals from CDKA;1-mVenus were detected with 514 nm excitation and 500-550 nm emission. Fluorescence confocal images were acquired using software cellSens (Evident, Tokyo, Japan).

#### RNA-seq

RNA Isolation was performed using Arabidopsis Col-0 roots for control, stress (salt, osmotic, cold, and heat), and stress recovery conditions using Qiagen RNeasy Plant Mini Kit (Cat.# 74904). The extracted RNA concentration and quality was analyzed using Thermo Scientific NanoDrop One (Cat.# 13-400-518). A total of 48 RNA samples (4 stresses X 3 conditions X 3 biological replicates) were considered for the experiment. Poly-A enriched RNA-seq libraries were prepared by using Vazyme VAHTS Universal V10 RNA-seq Library Prep Kit and sequenced at 150-bp PE on an Element AVITI instrument by AmpSeq (Gaithersburg, Maryland, USA) (https://www.ampseq.com/).

Reads were filtered with fastp v0.23.1 (*50*), mapped with STAR v2.7.11 to the TAIR10 Arabidopsis genome (*51*, *52*), and quantified with Salmon v1.10.2 (*53*). Four samples were outliers and excluded from analysis. Differential gene expression was called at FDR<0.05 with package DESeq2 v1.44.0 in R (*54*, *55*) and visualized with package pheatmap v1.0.12 (*56*).

#### Analyzing single cell sequencing data

*CYCB1;1*, *CDKA;1*, and *ICK1* gene expression was visualized using Trevor Nolan’s lab (https://shiny.mdc-berlin.de/ARVEX/) and Root Cell Atlas (https://rootcellatlas.org/) visualization tools for already published single cell sequencing data of Arabidopsis root.

#### qPCR

Each RNA concentration was normalized with RNase-free water, and 500 ng RNA was used to synthesize cDNA using NEB ProtoScript II Reverse Transcriptase (Cat.# M0368S). Quantitative PCR reactions were performed using StepOnePlus Real-Time PCR System (Applied Biosystems) and PowerUp SYBR Green Master Mix (Applied Biosystems; Cat.# A25741). The reaction was performed as per the manufacturer’s instructions. The list of primers used in this study is mentioned in Supplemental Table 3.

#### Molecular docking and simulation

Structural models of CDKA1 bound to ATP and olomoucine were predicted using the AlphaFold Server (*30*) and DynamicBind (*31*) via the Neurosnap Inc web servers (https://neurosnap.ai/). Molecular dynamics simulations were conducted using GROMACS (*32*). Parameters included the AMBER99SB-ILDN force field, a cubic solvent box with an ionic concentration of 0.15 M over a 1 nanosecond simulation time. All simulations were carried out using the Neurosnap Inc server (https://neurosnap.ai/). Simulation H-bond occupancies were analyzed using Galaxy Cheminformatics web servers, with an angle cutoff of 20 degrees and distance cutoff of 3 Ångstroms (*57*, *58*). Molecular visualization and structural rendering of CDKA1 and ligands were performed using ChimeraX 1.4 (*33*).

## Statistical analysis

The raw data for each quantification was imported to statistical software JMP Pro17 for generating graphs and performing statistical tests.

## Acknowledgements

The authors thank Masaaki Umeda (Nara Institute of Science and Technology), Sachihiro Matsunaga (Tokyo University of Science), Arp Schnittger (University of Hamburg), and Hong Wang (University of Saskatchewan) for sharing Cytrap (*17*), *pAtPCNA1::AtPCNA1-sGFP* (*19*), CDKA;1-mVenus/*cdka;1* (*48*), and *35S::GFP-ICK1* (*42*), respectively, seeds. Authors thank Miki Fujita and EunKyoung Lee of UBC Bioimaging Facility (RRID: SCR_021304) for their kind support.

## Funding

The research at Ashraf lab is funded by the NSERC Discovery grant (RGPIN-2025-04277) and start-up grant provided by the Department of Botany and Faculty of Science at the University of British Columbia. Olivia S. Hazelwood is supported by the 4 years doctoral fellowship program by the University of British Columbia. Research in Gallagher and Herb labs is supported by the US Department of Agriculture Agricultural Research Service (2072-21500-001-000D).

## Authors contributions

O.S.H., K.A.D., V.H., D.H., J.P.G., M.A.A. designed and performed experiments and analyzed data. O.S.H., J.P.G., and M.A.A. wrote the manuscript and all the authors agreed with the final version of the manuscript.

## Competing interests

Authors declare no conflict of interests.

## Data and materials availability

RNA-seq data are available on NCBI SRA under BioProject PRJNA1253770.

**Supplemental Table 1:** Read counts for RNA-seq libraries.

**Supplemental Table 2:** Differentially expressed genes in RNA-seq experiment.

**Supplemental Table 3:** List of primers used in this study for qPCR experiment.

**Supplemental movie 1:** Molecular dynamic simulation of CDKA;1 (cyan) and olomoucine (red). The interacting amino acid residues are highlighted in yellow.

**Supplemental movie 2:** Molecular dynamic simulation of CDKA;1 (cyan) and ATP (magenta). The interacting amino acid residues are highlighted in yellow.

---

**Supplemental figure 1:**
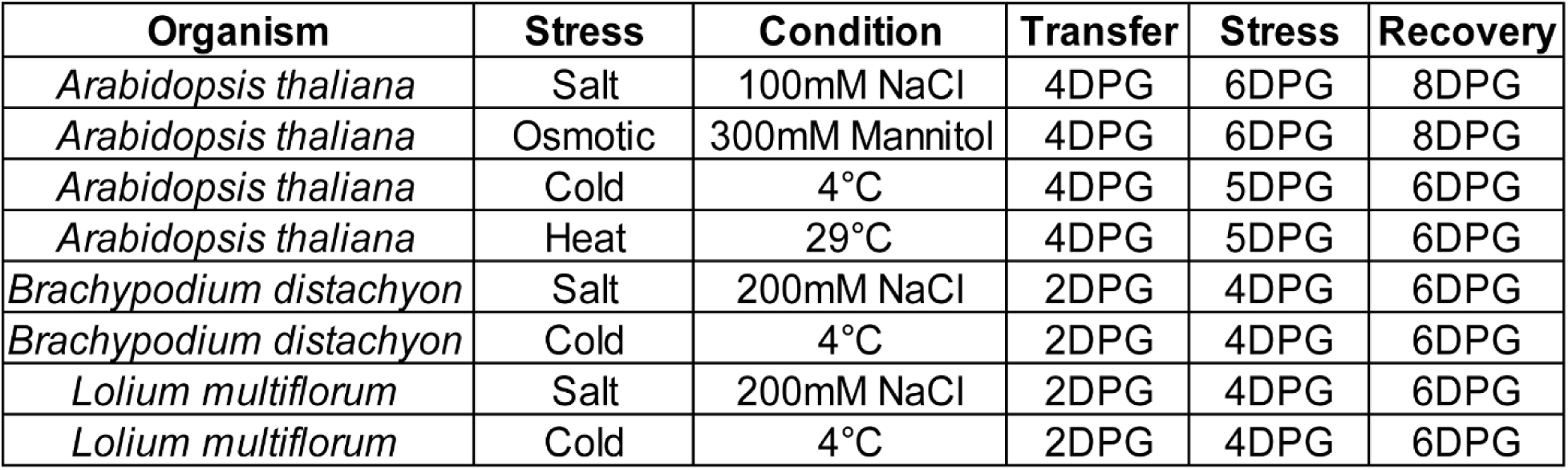
Experimental conditions for Arabidopsis, Brachypodium, and annual ryegrass. Concentration of salt and mannitol, temperature for cold and heat stress, and developmental stage of root, and incubation periods were mentioned for each individual stress and each organism used in this study.

**Supplemental figure 2:**
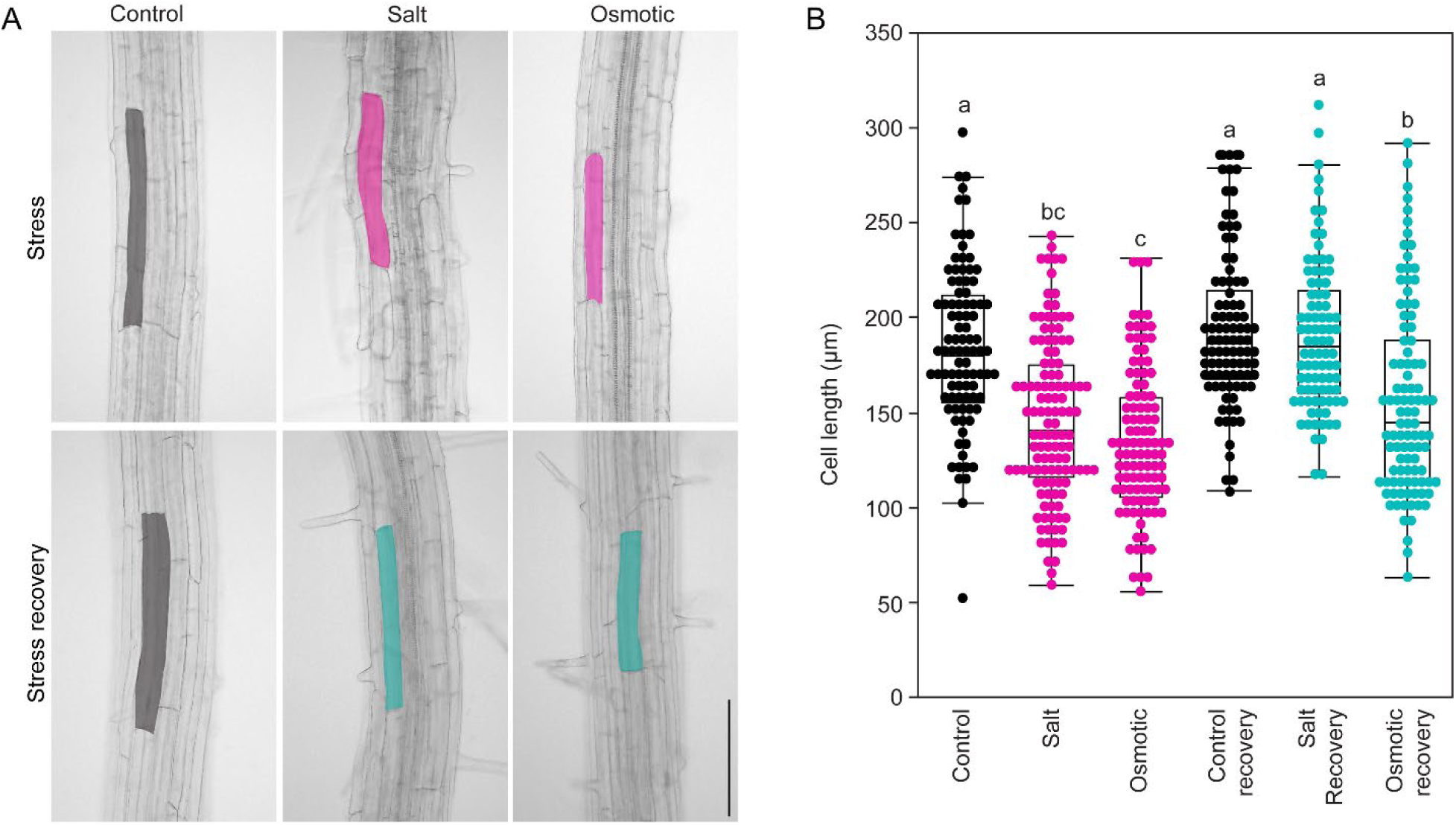
Cell elongation during salt and osmotic stress, and stress recovery period. (A) Epidermal cell length was measured during the control, salt and osmotic stress, and corresponding recovery period. Scale bar = 100 µm. (B) Quantification of cell length from A. Statistical test is performed based on Tukey’s Honest test. Groups labeled with the same letter are not statistically different from each other (alpha = 0.05). Boxplots show median values (center line), 25th to 75th interquartile range (box) and 1.5*interquartile range (whiskers).

**Supplemental figure 3:**
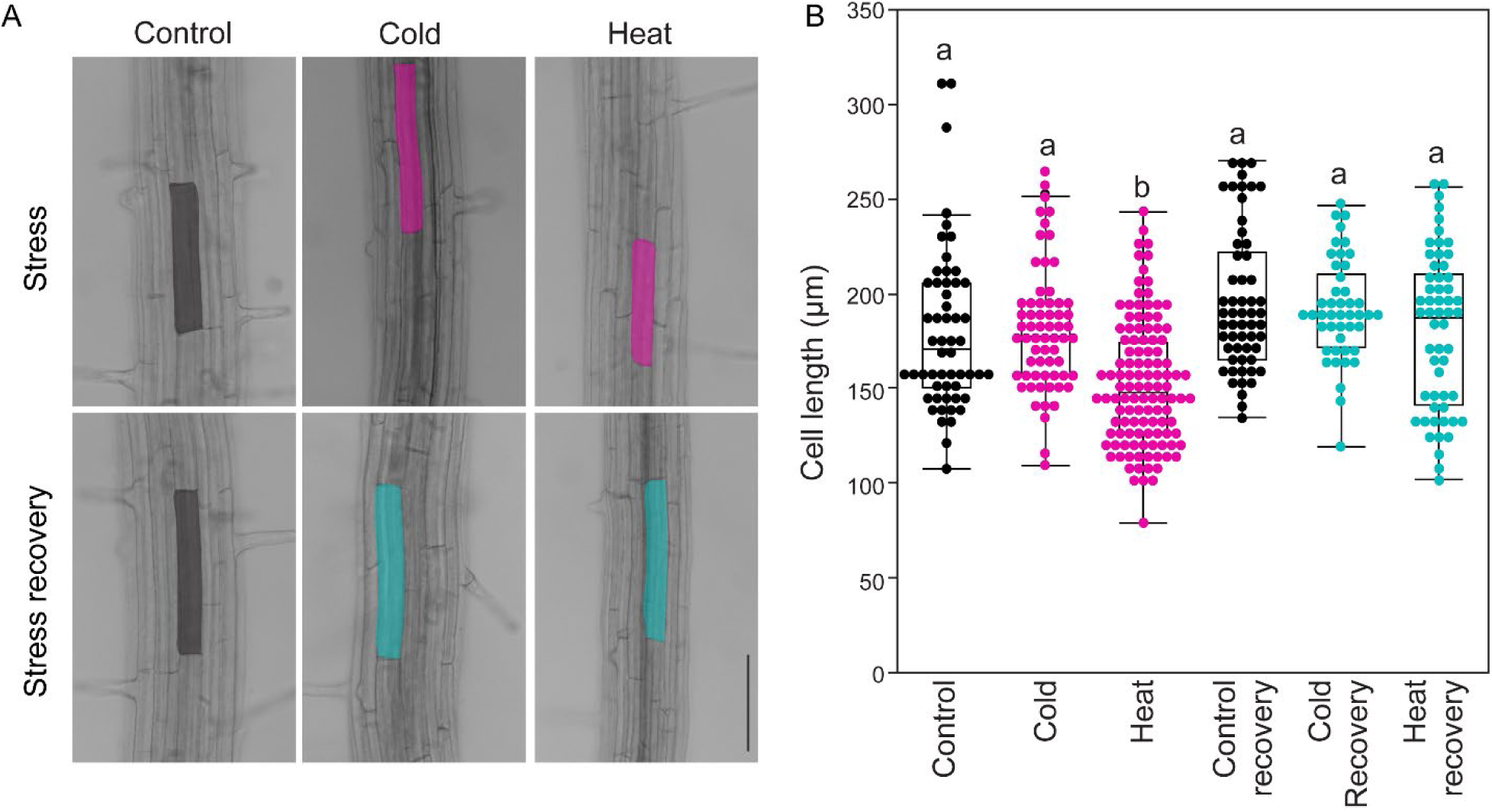
Cell elongation during cold and heat stress, and stress recovery period. (A) Epidermal cell length was measured during the control, cold and heat stress, and corresponding recovery period. Scale bar = 100 µm. (B) Quantification of cell length from A. Statistical test is performed based on Tukey’s Honest test. Groups labeled with the same letter are not statistically different from each other (alpha = 0.05). Boxplots show median values (center line), 25th to 75th interquartile range (box) and 1.5*interquartile range (whiskers).

**Supplemental figure 4:**
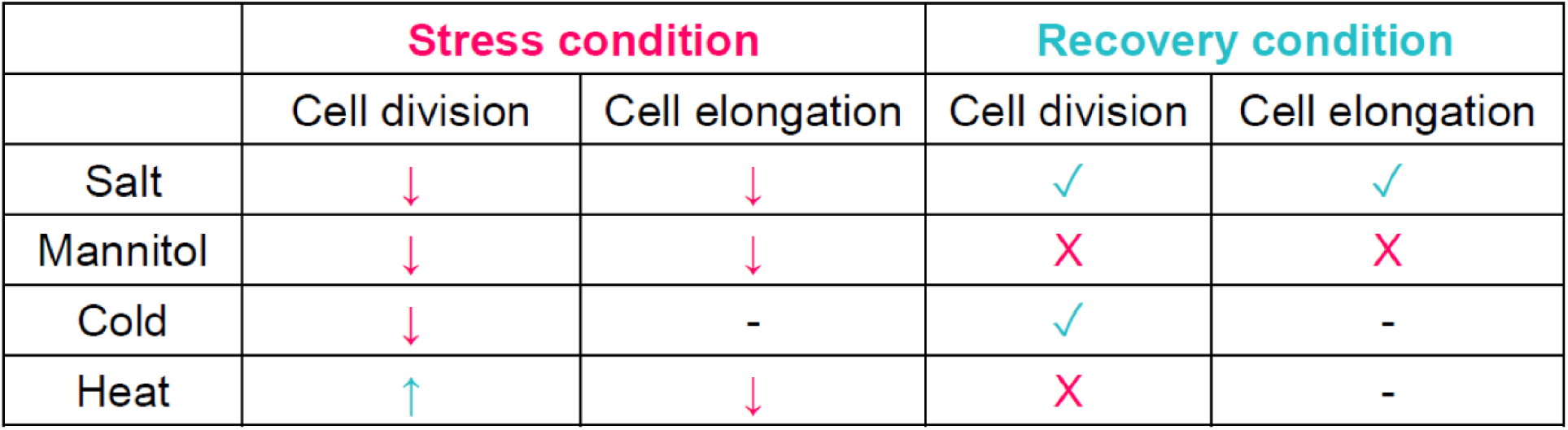
Summary of cell division and cell elongation during salt, osmotic, cold, and heat stress, and stress recovery period. The summary is based on Figure 1, 2, and Supplemental figure 2, 3. Upward and downward arrow during stress condition indicates increased cell division or cell elongation. Dash line indicates neutral or wild-type response to the stress. Tick and cross marks indicate recovered and not recovered cell division or cell elongation during the recovery period. Dash line indicates neutral or wild-type response during the recovery period.

**Supplemental figure 5:**
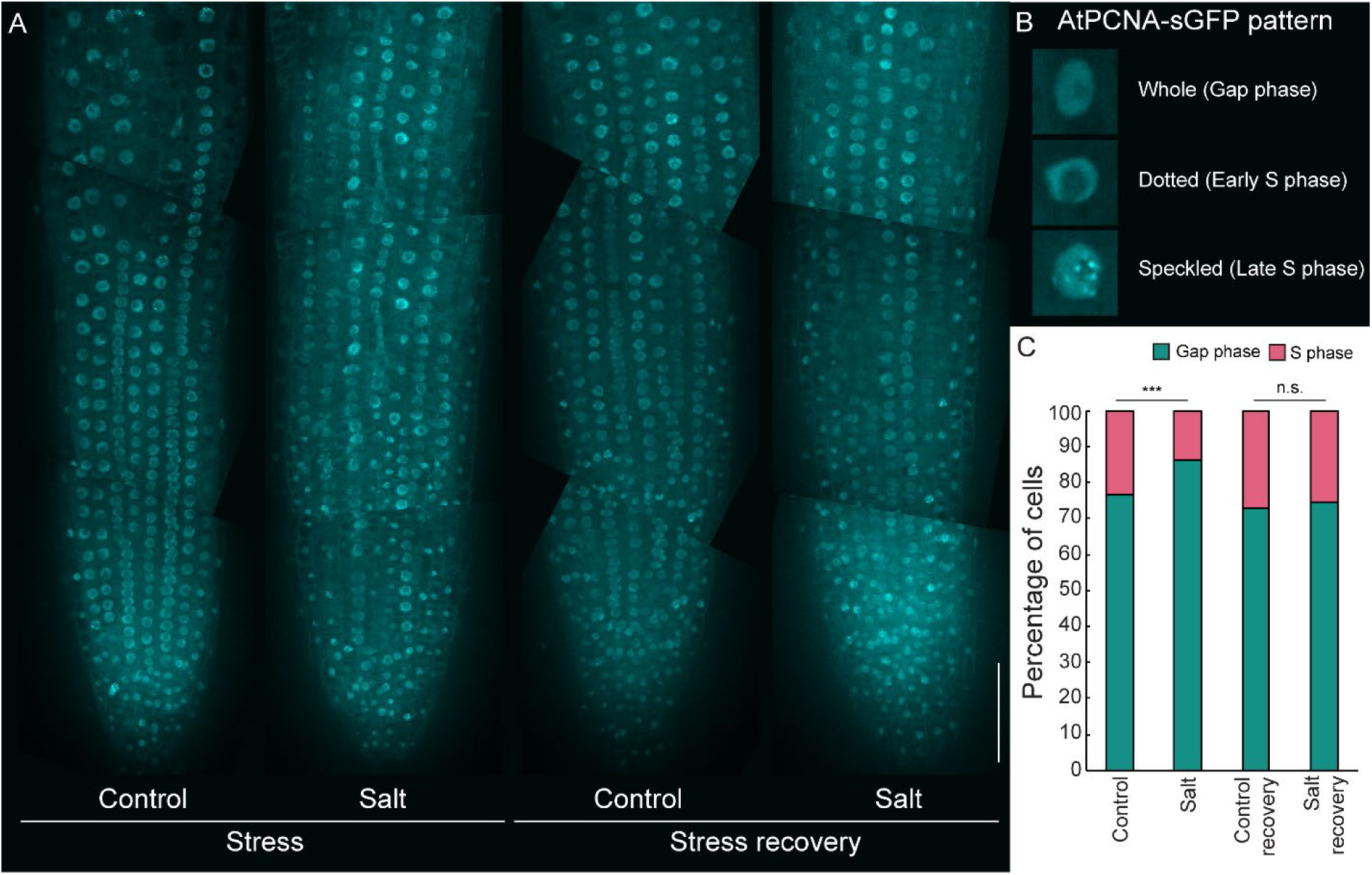
PCNA-GFP signal during salt stress and recovery. (A) Cell cycle marker PCNA-GFP signal during salt stress and recovery. Scale bar = 100 µm. (B) Characteristic pattern of whole (Gap phase), dotted (early S phase), and speckled (late S phase) pattern of PCNA-GFP marker. (C) Quantification of number of cells in gap phase and S phase during salt stress and recovery. Statistical test was performed based on Fisher’s exact test.

**Supplemental figure 6:**
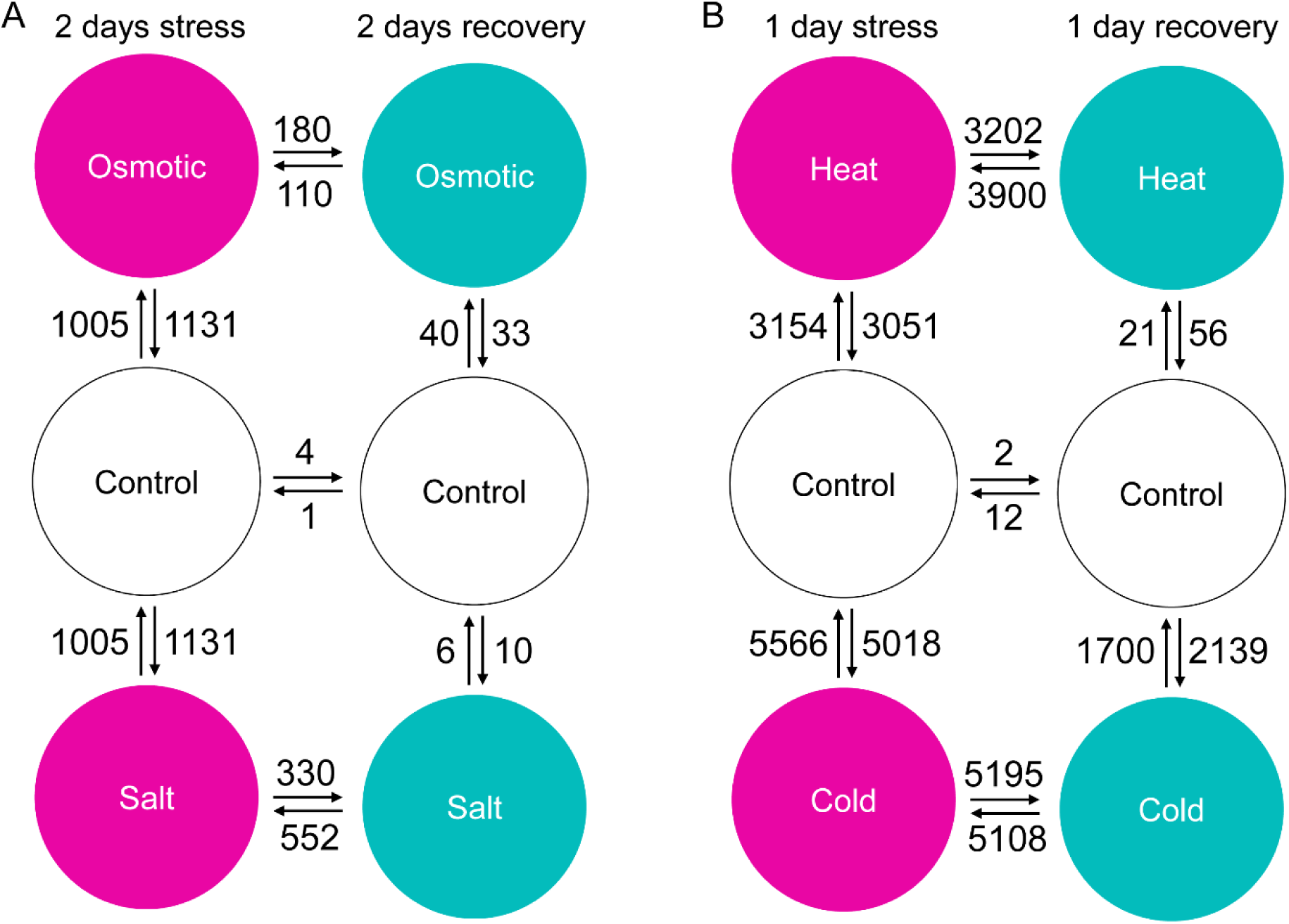
Differential gene expression from comparative transcriptomics. Number of differentially expressed genes during stress [salt (A), osmotic (A), cold (B), and heat (B)] and stress recovery are highlighted.

**Supplemental figure 7:**
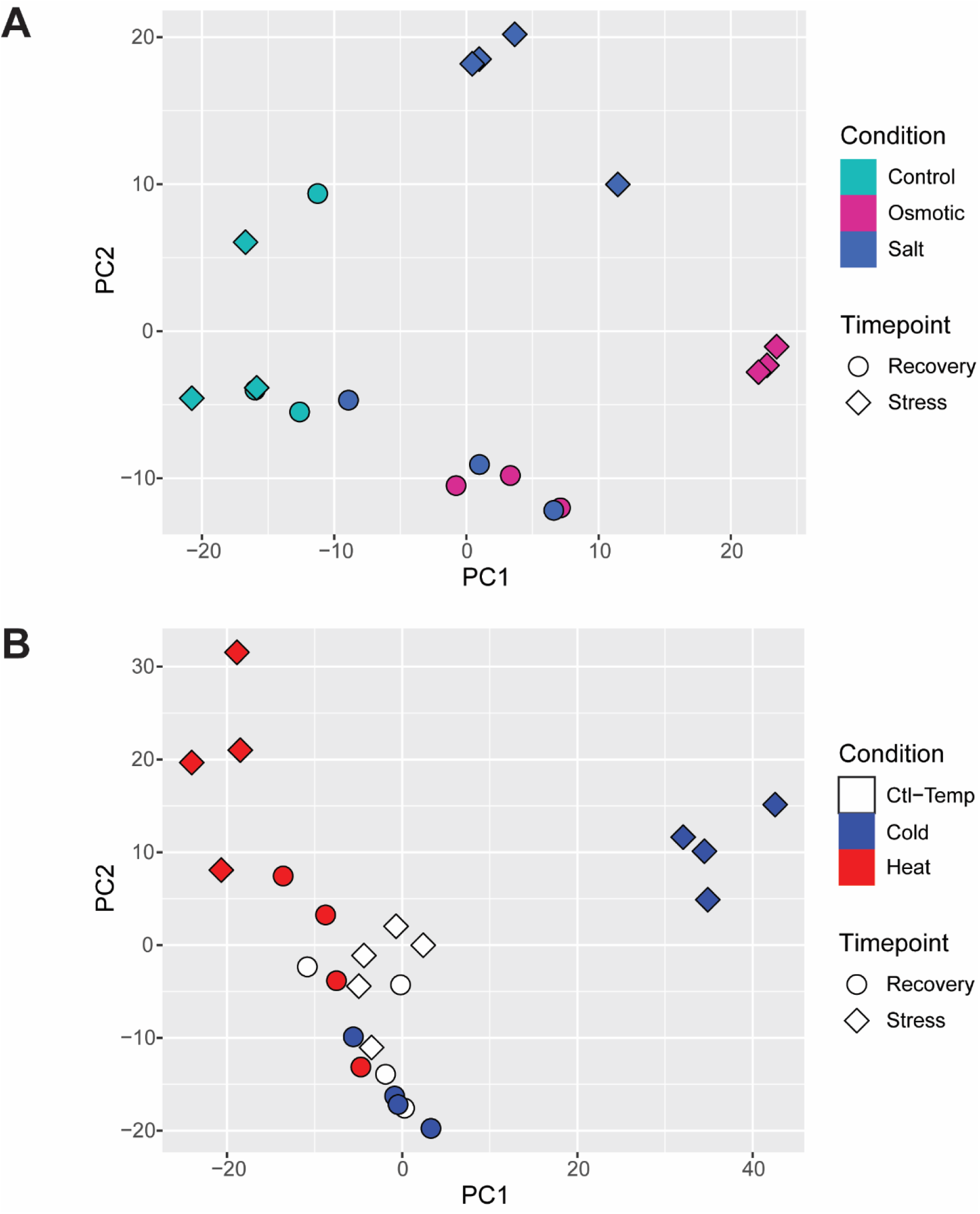
Principal component analysis (PCA) of RNAseq samples. PCA demonstrates the clustering of similar RNAseq samples for control, stress, control recovery, and stress recovery in A (salt and osmotic stress) and B (cold and heat stress).

**Supplemental figure 8:**
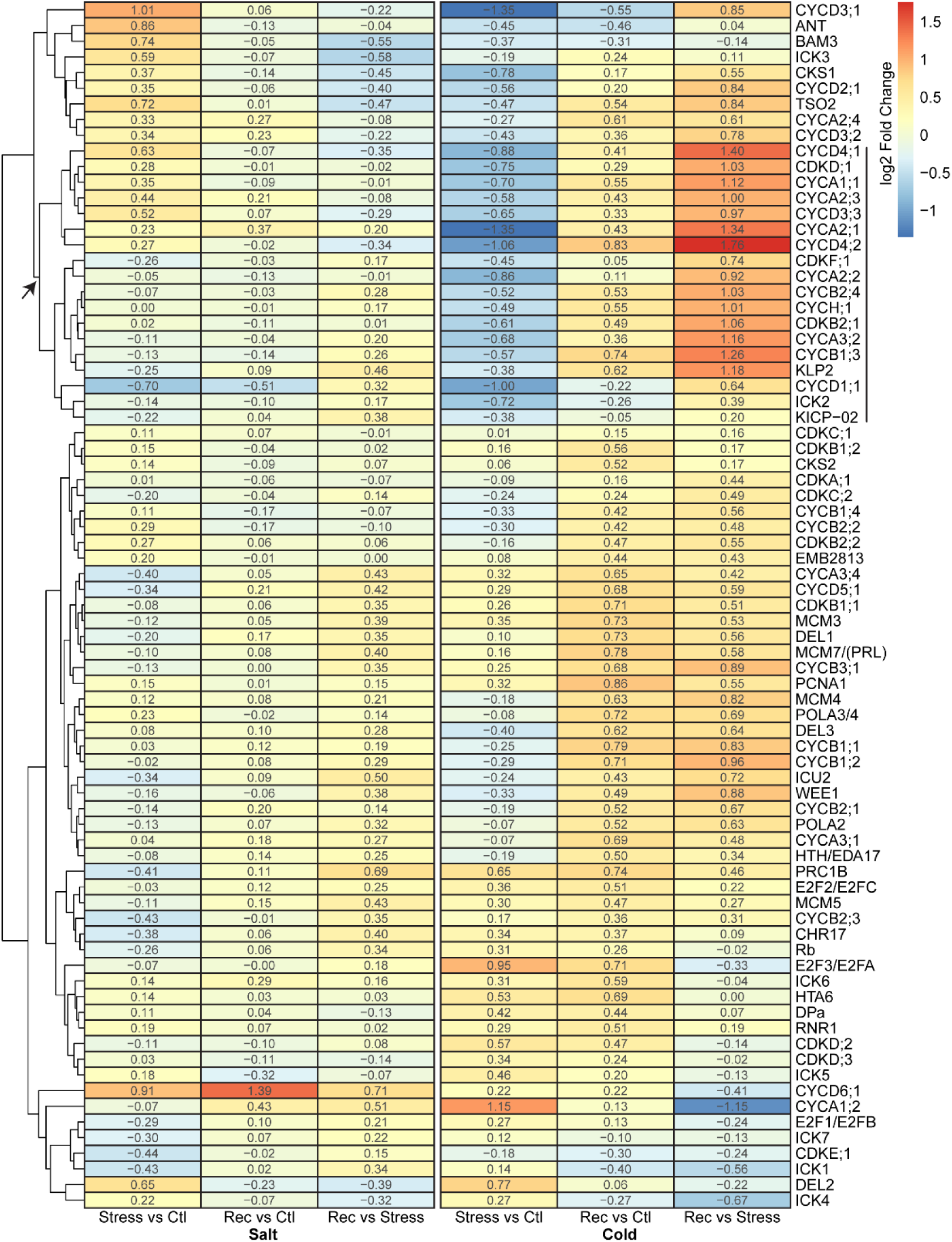
Differential expression of core cell cycle genes in Arabidopsis. Log2Fold changes of core cell cycle genes based on the comparative transcriptomics during salt and cold stress, and stress recovery. The gene clustering highlights a subset of genes used in Figure 3A.

**Supplemental figure 9:**
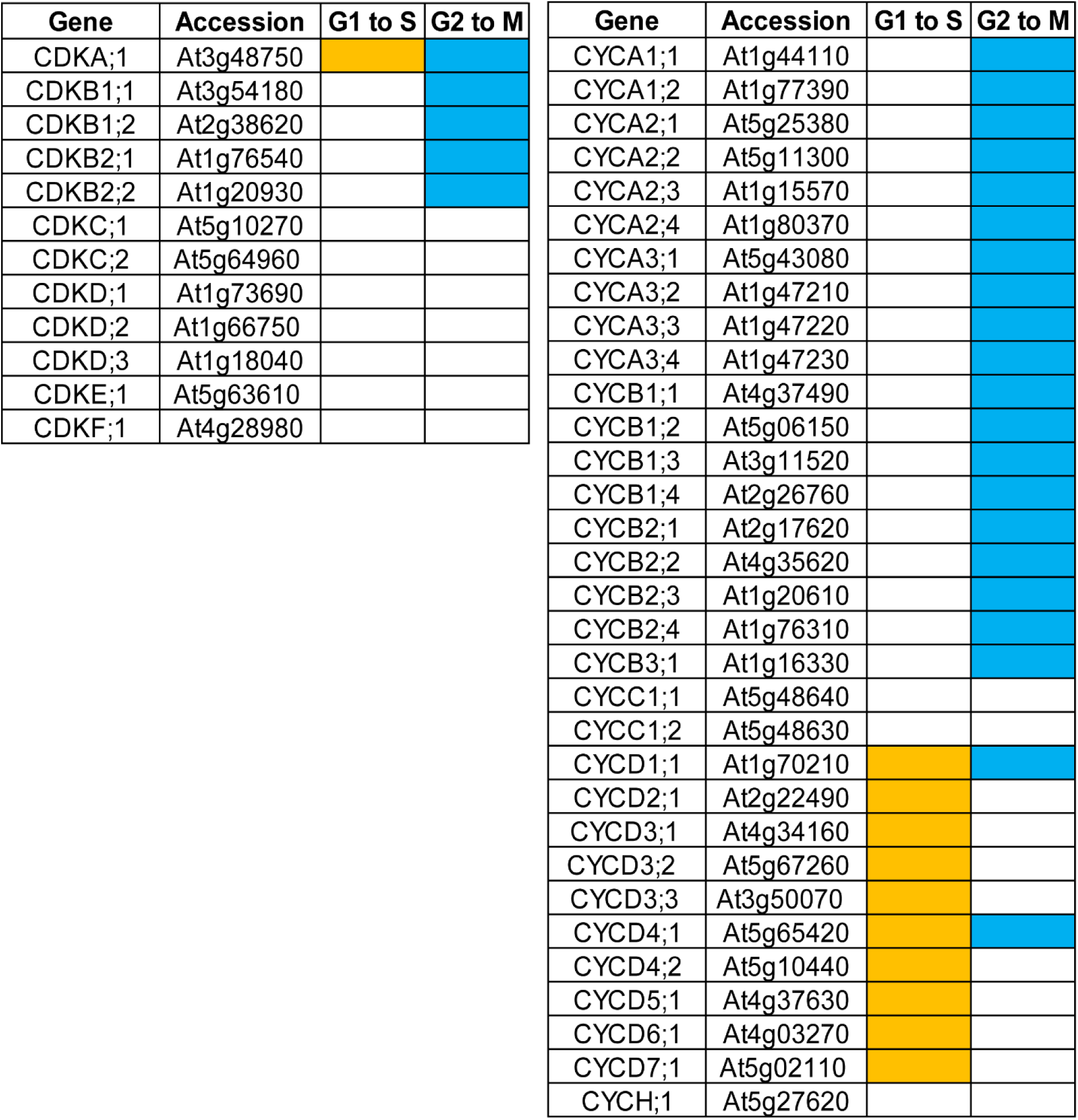
Cell cycle specific function of CDKs and cyclins. List of Arabidopsis CDKs and cyclins involved in G1 to S (yellow) and G2 to M (cyan) transitions.

**Supplemental figure 10:**
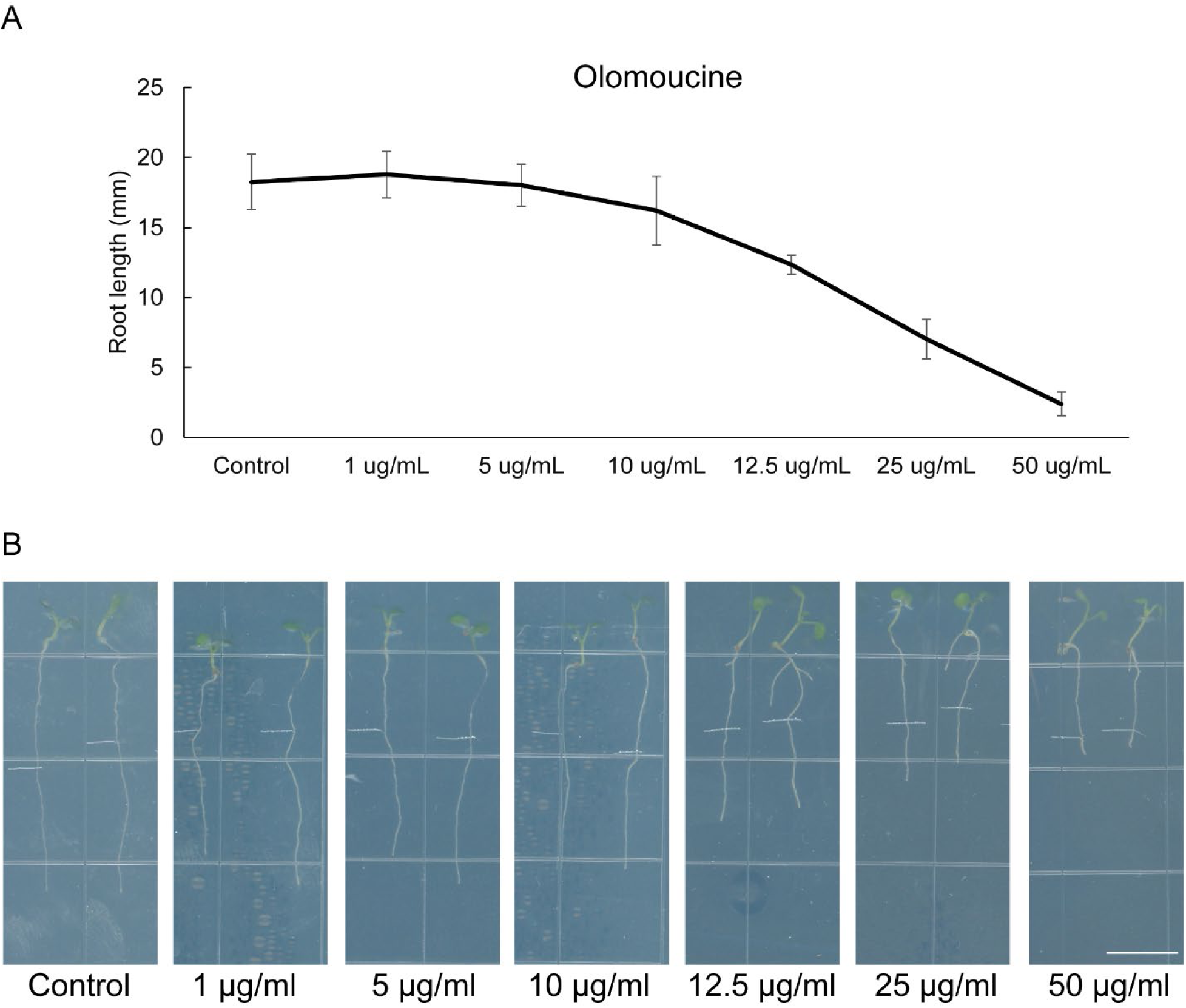
Dose response curve of olomoucine. (A) Quantification of root length in presence of 1, 5, 10, 12.5, 25, and 50 µg/ml olomoucine. (B) Representative images for each olomoucine concentration corresponding to A. Scale bar = 10 mm.

**Supplemental figure 11:**
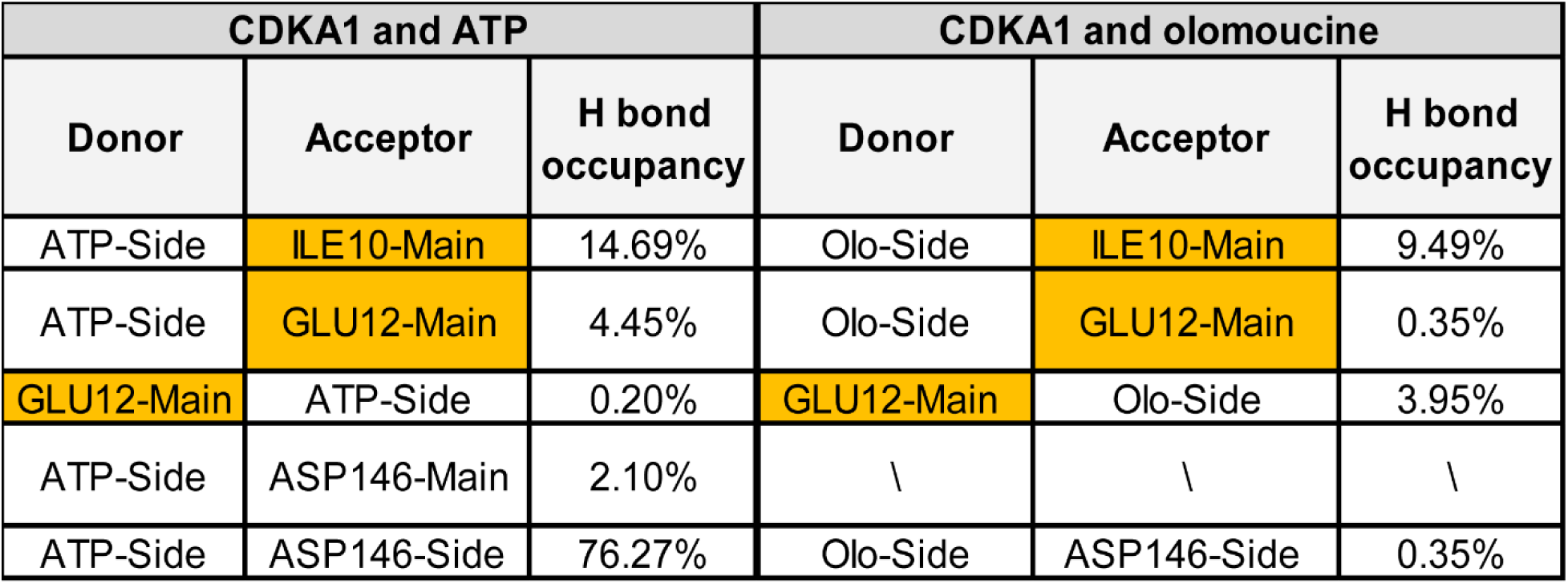
Hydrogen bond occupancy of ATP and olomoucine with CDKA1. Interacting amino acids responsible for maintaining interaction with CDKA1 for both ATP and olomoucine; and hydrogen bond occupancy for each interaction.

**Supplemental figure 12:**
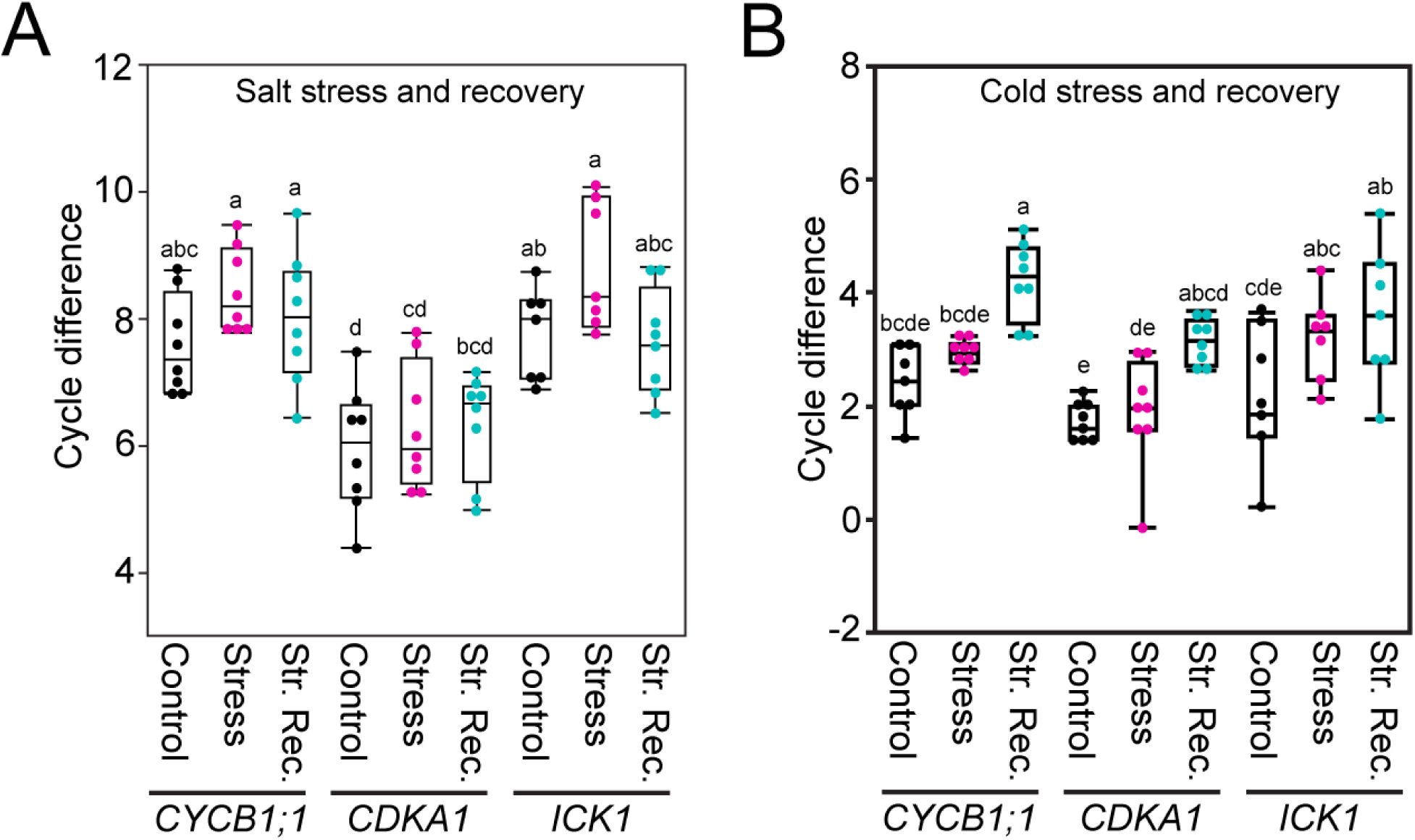
Real time PCR of *CYCB1;1* (left), *CDKA1* (middle), and *ICK1* (right) genes in salt (A) and cold (B) stress, and corresponding recovery periods. Statistical test is performed based on Tukey’s Honest test (A and B). Groups labeled with the same letter are not statistically different from each other (alpha = 0.05). Boxplots show median values (center line), 25th to 75th interquartile range (box) and 1.5*interquartile range (whiskers) (A and B).

## Notes

### Competing Interest Statement

The authors have declared no competing interest.

### Summary of Updates

We have updated the Figure 3 and 4, created supplemental figure 12, and also the explanation in the text section based on the comments received from colleagues.

