## Supplementary figures and images for "Cell cycle follows “pause and play” mechanism in environment stress recovery in diverse plant species"

### Supplemental_Figure1.tif

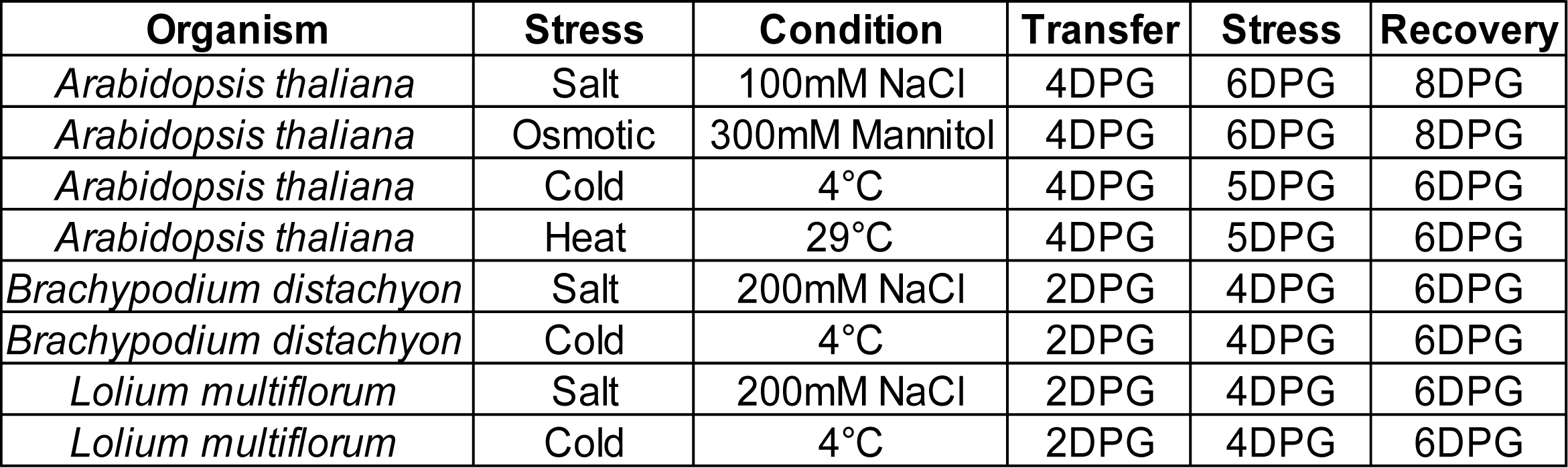

### Supplemental_Figure2.tif

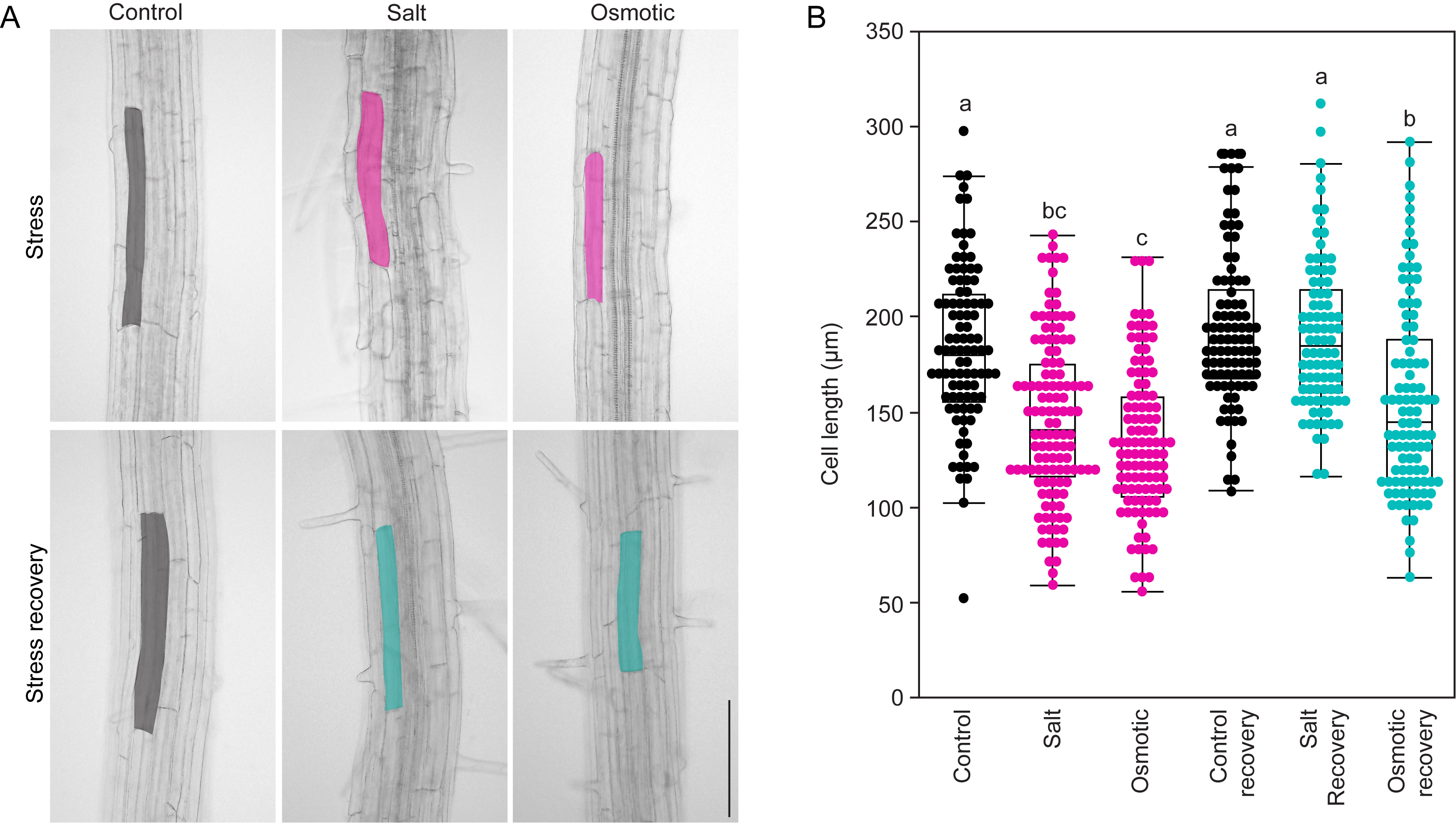

### Supplemental_Figure3.tif

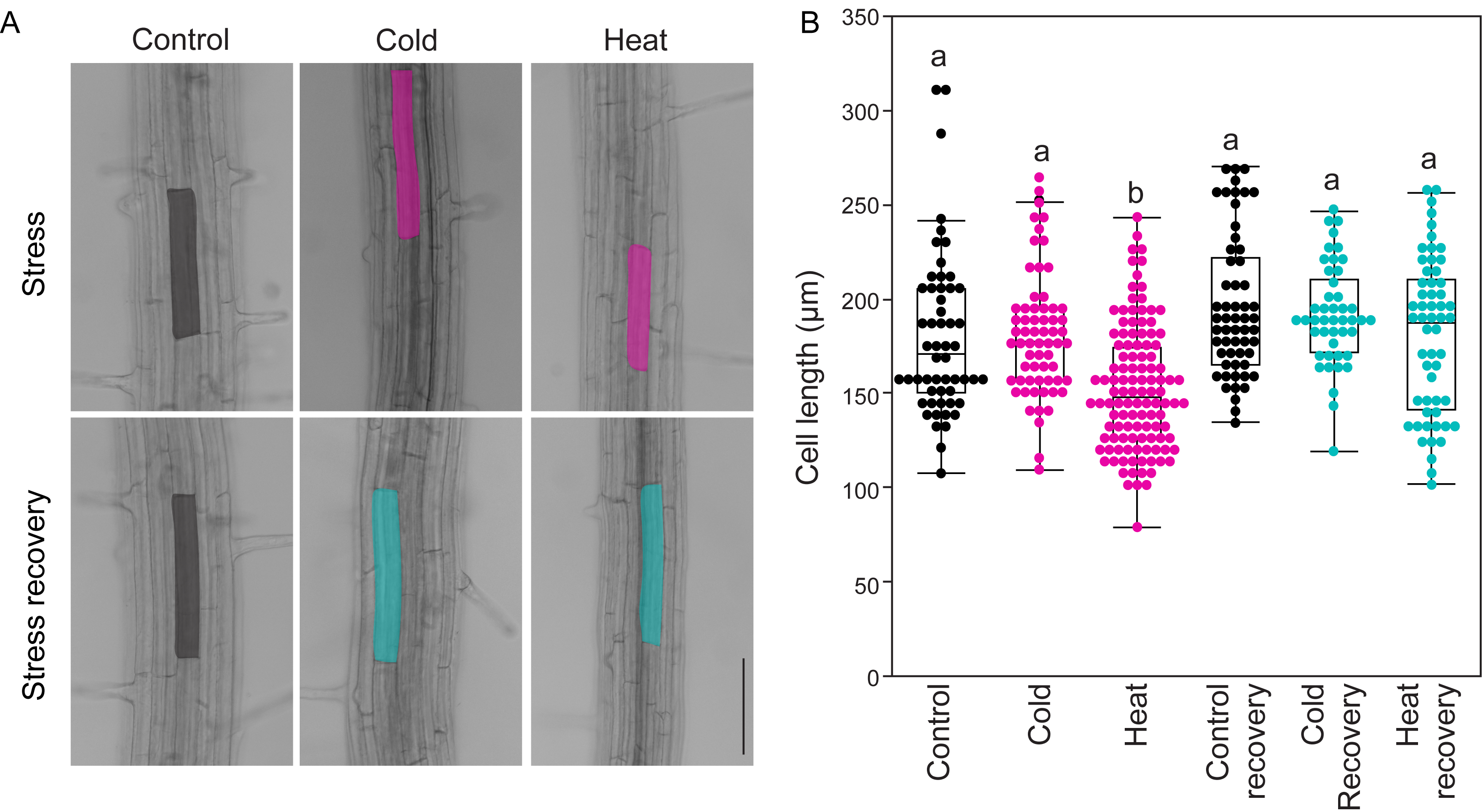

### Supplemental_Figure4.tif

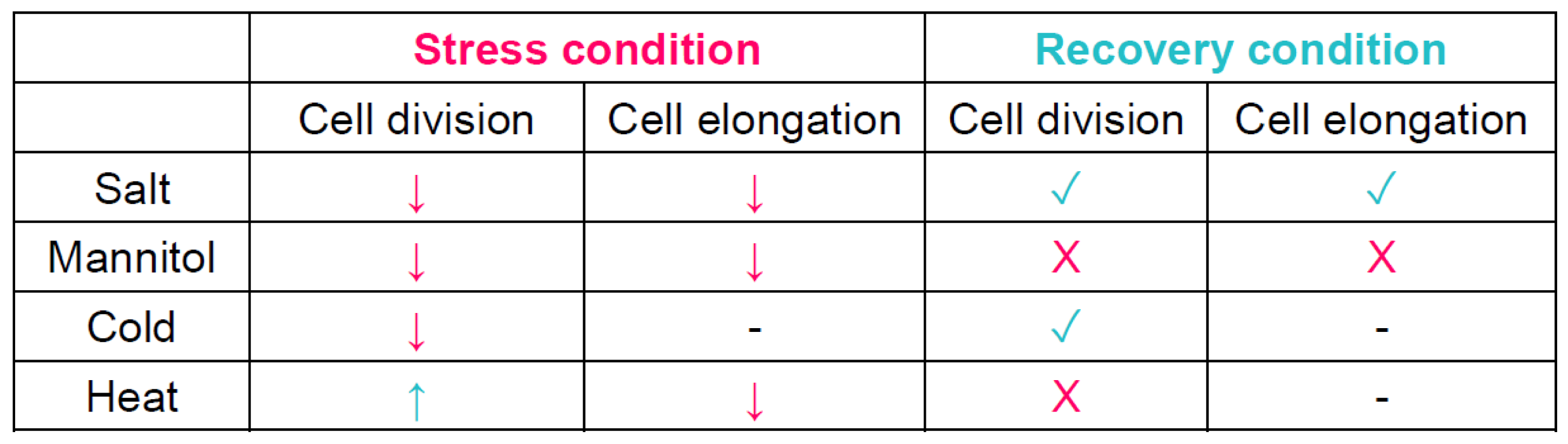

### Supplemental_Figure5.tif

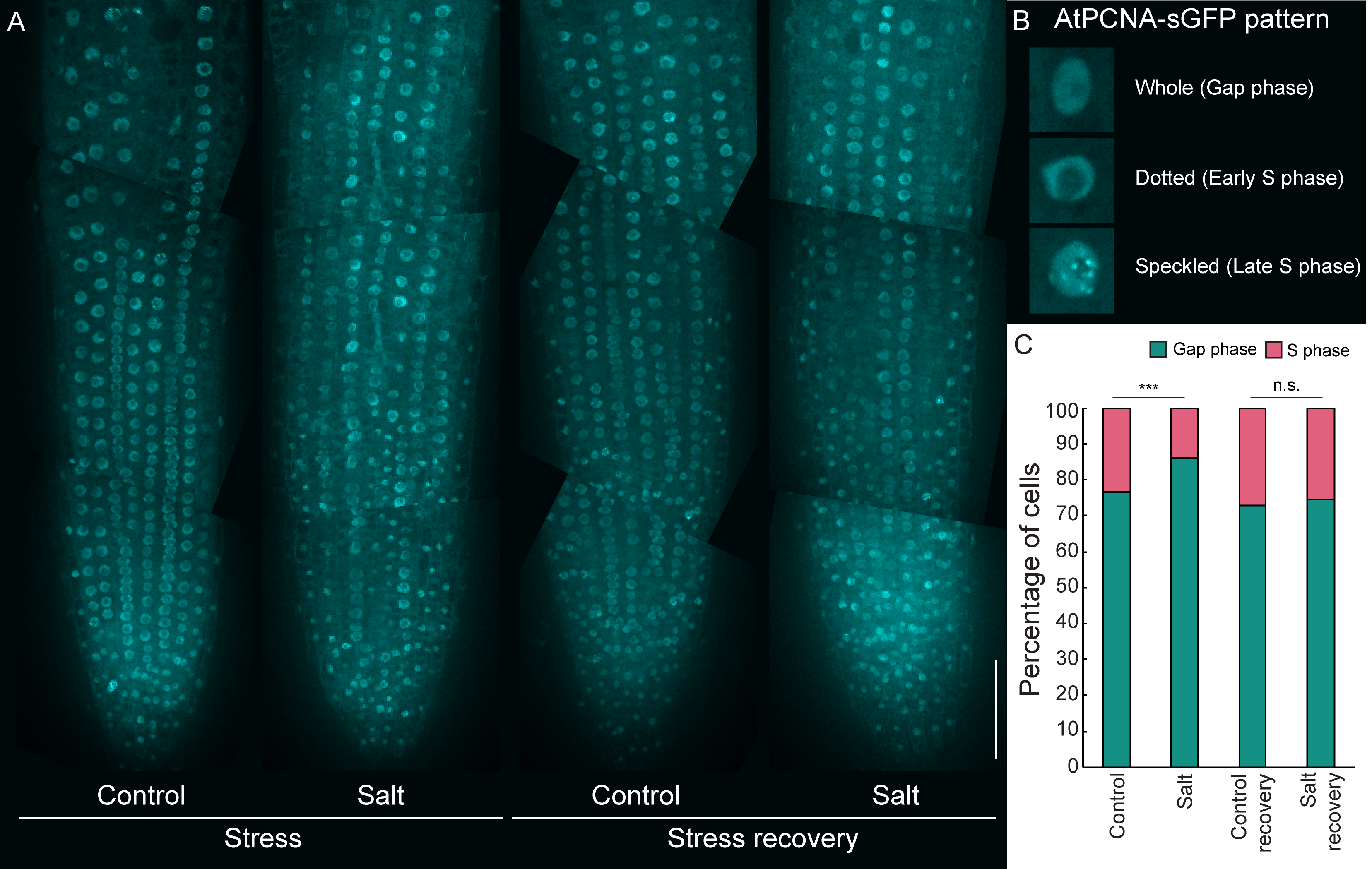

### Supplemental_Figure6.tif

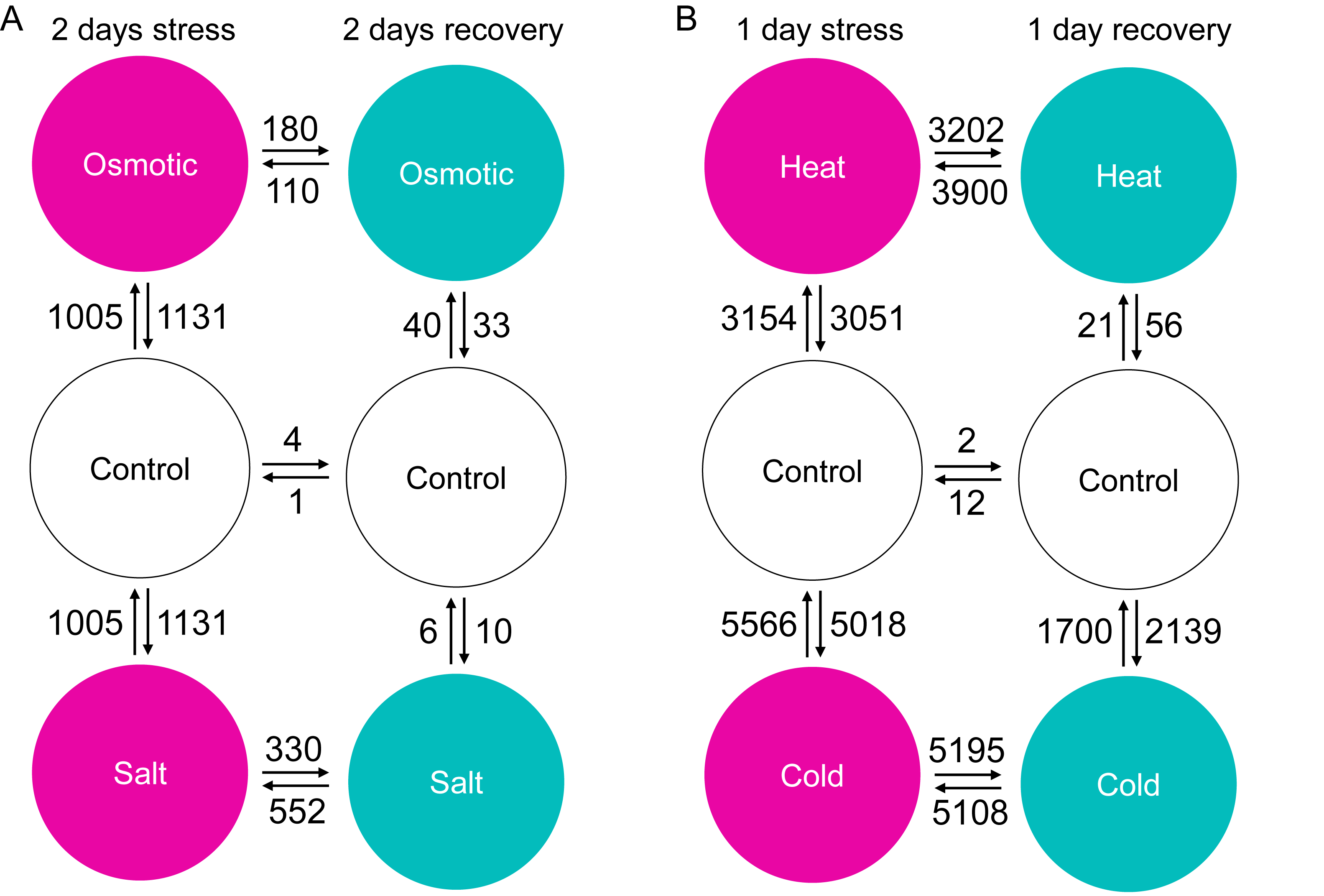

### Supplemental_figure7.tif

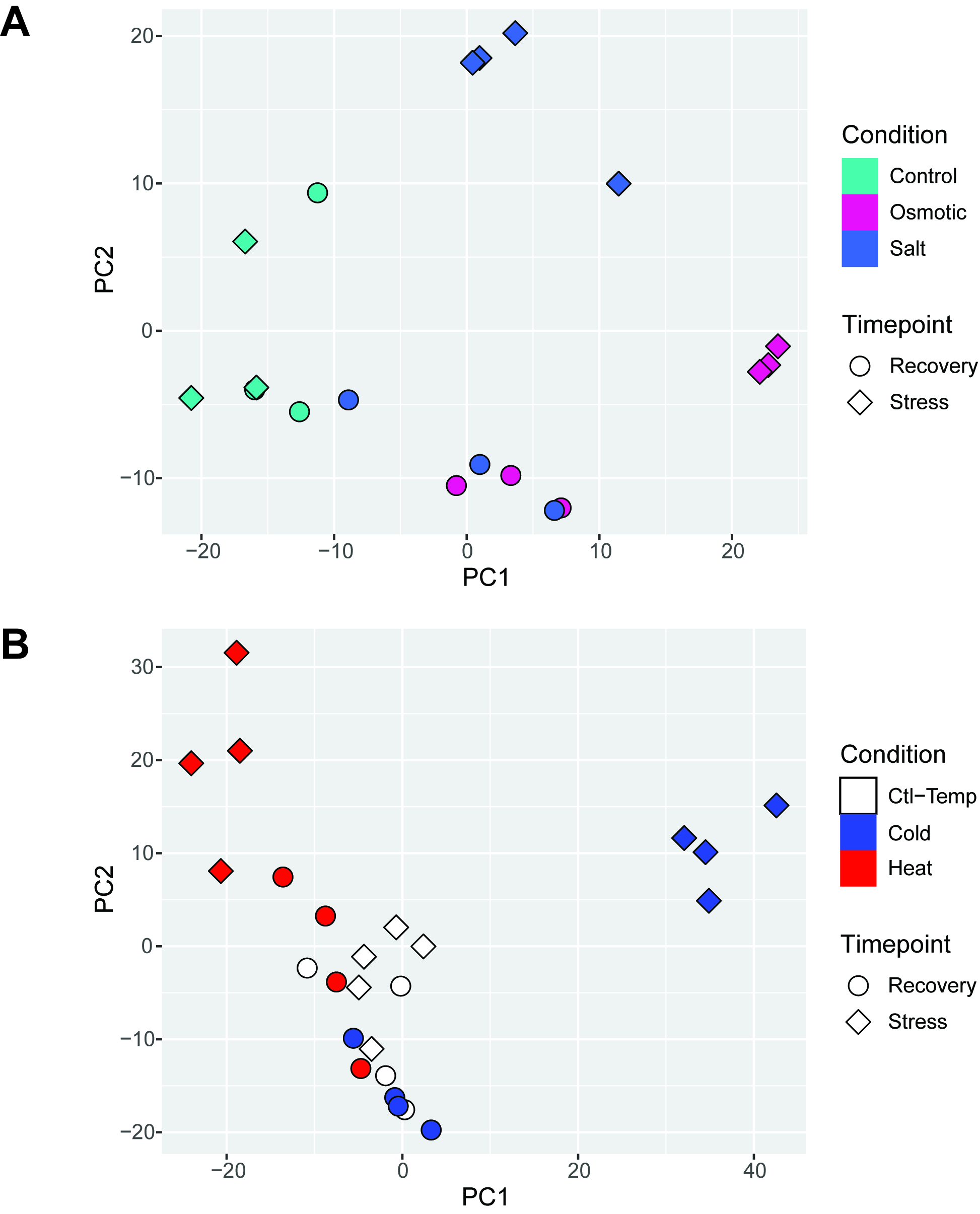

### Supplemental_Figure8.tif

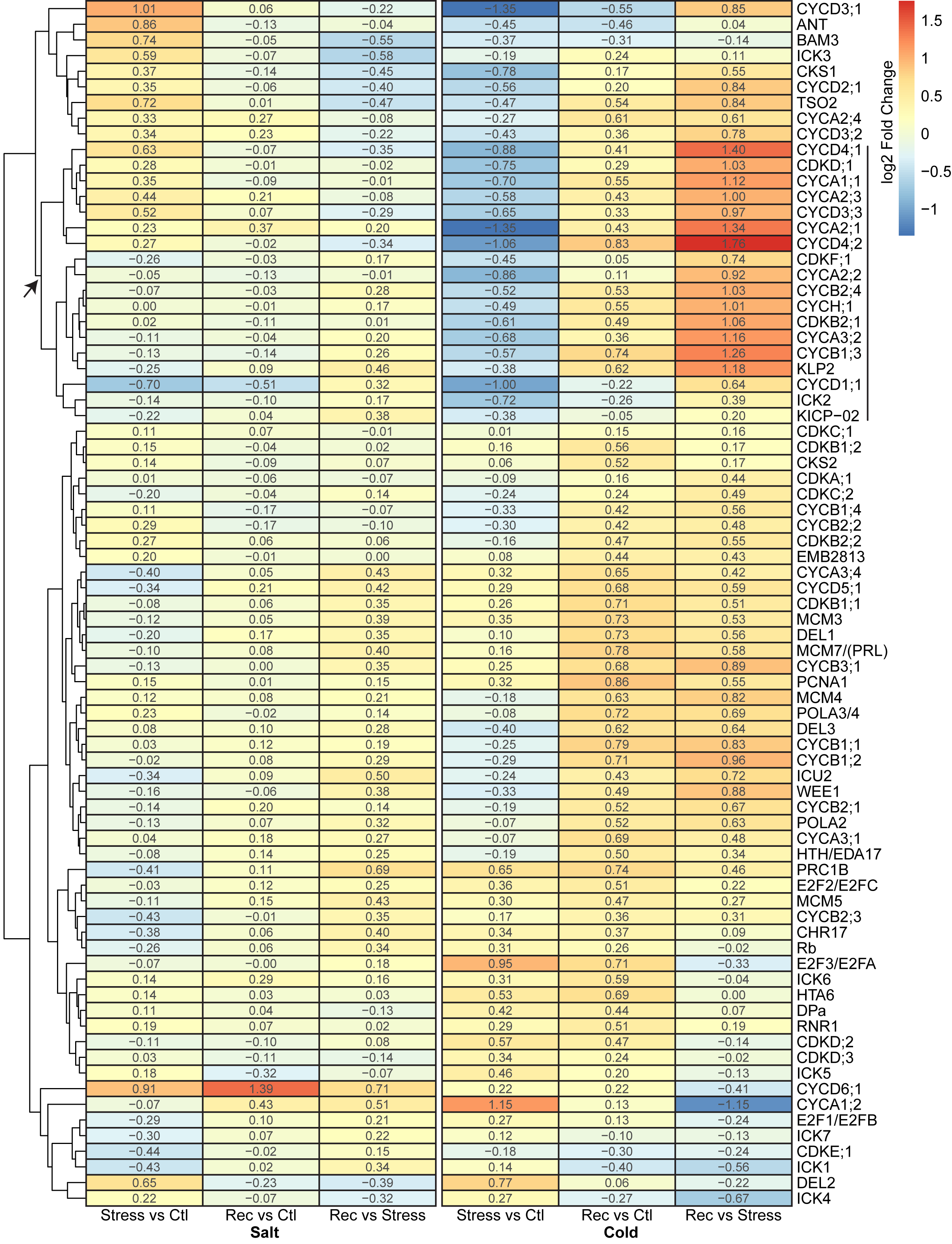

### Supplemental_Figure9.tif

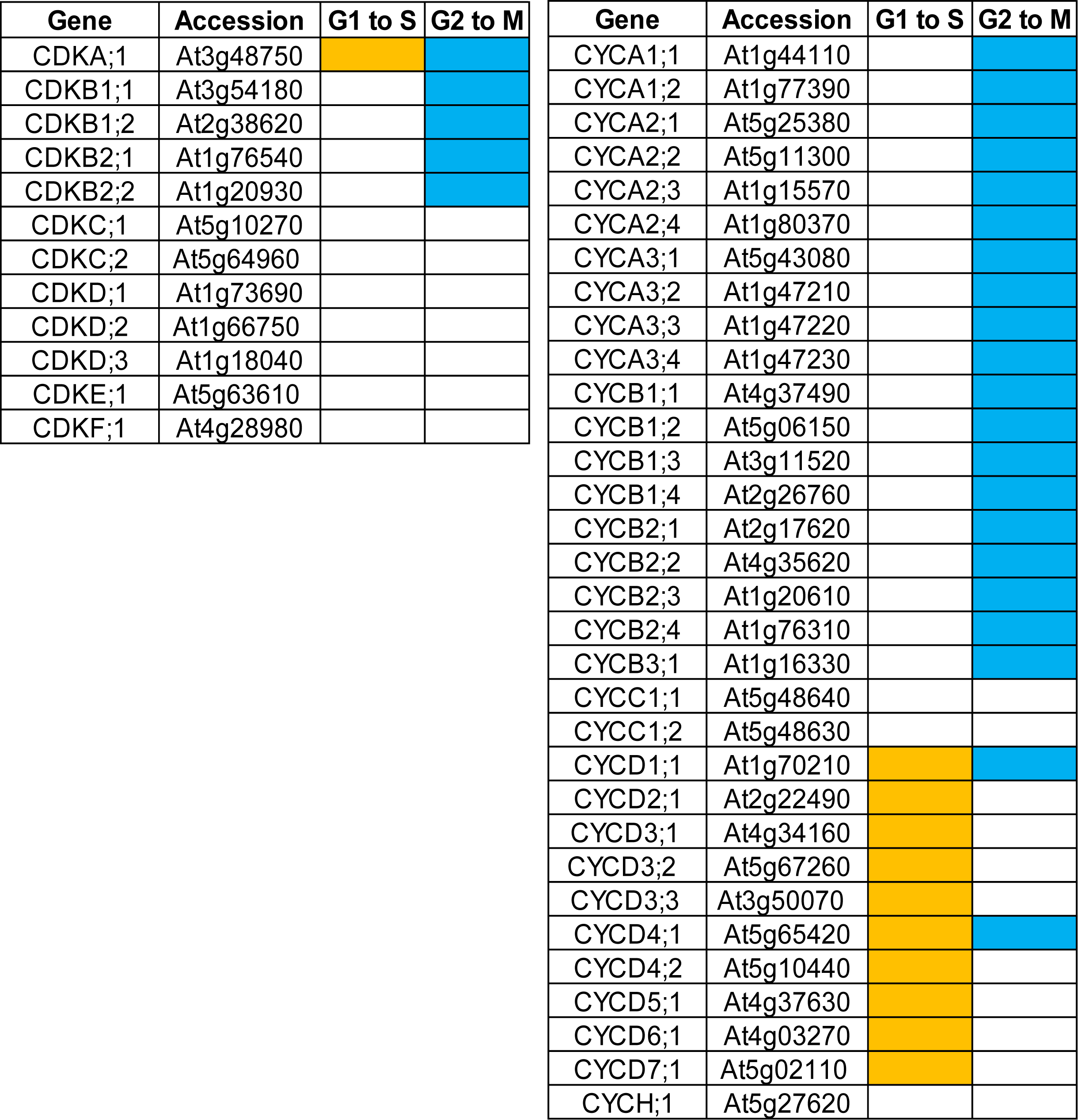

### Supplemental_Figure10.tif

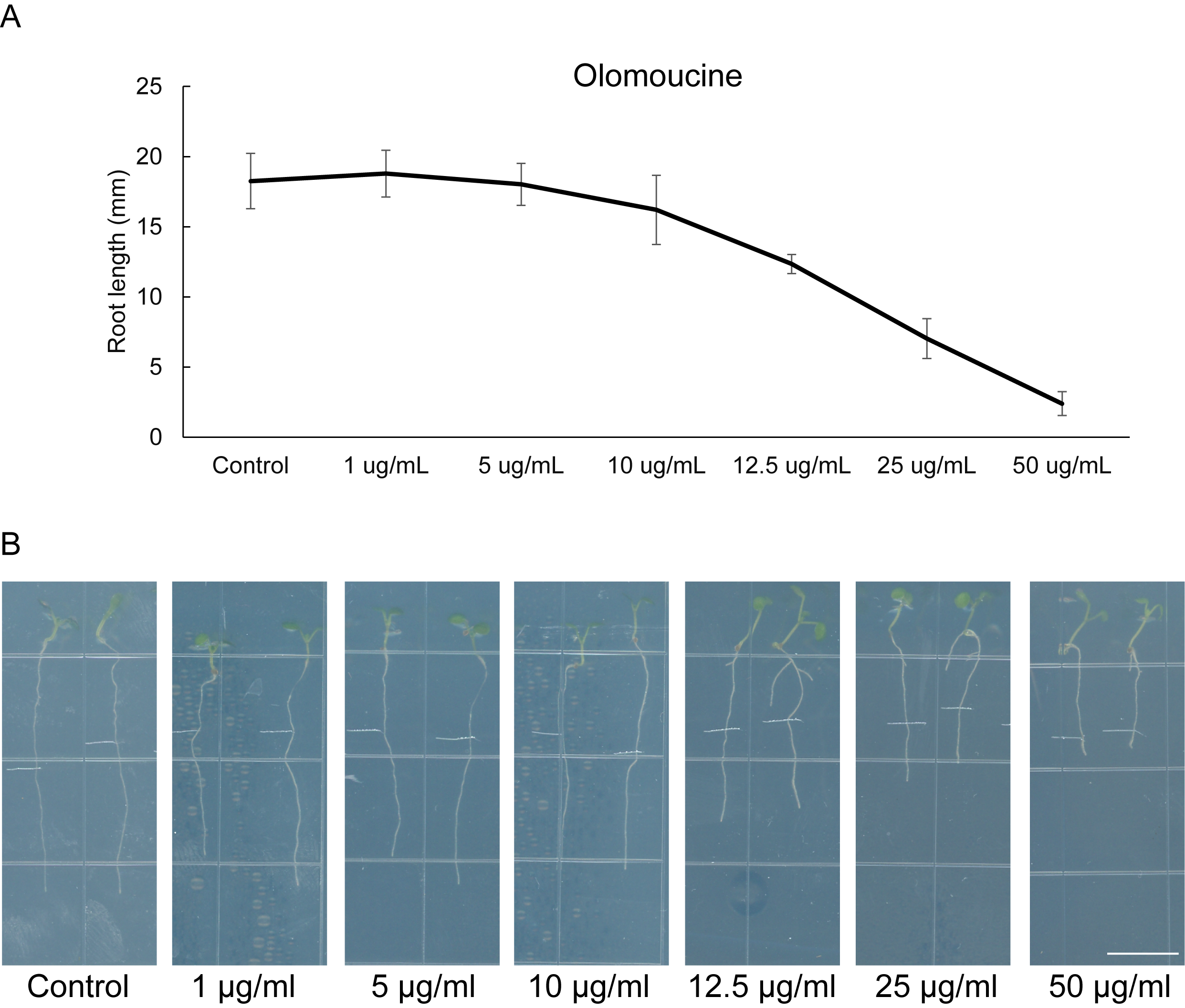

### Supplemental_Figure11.tif

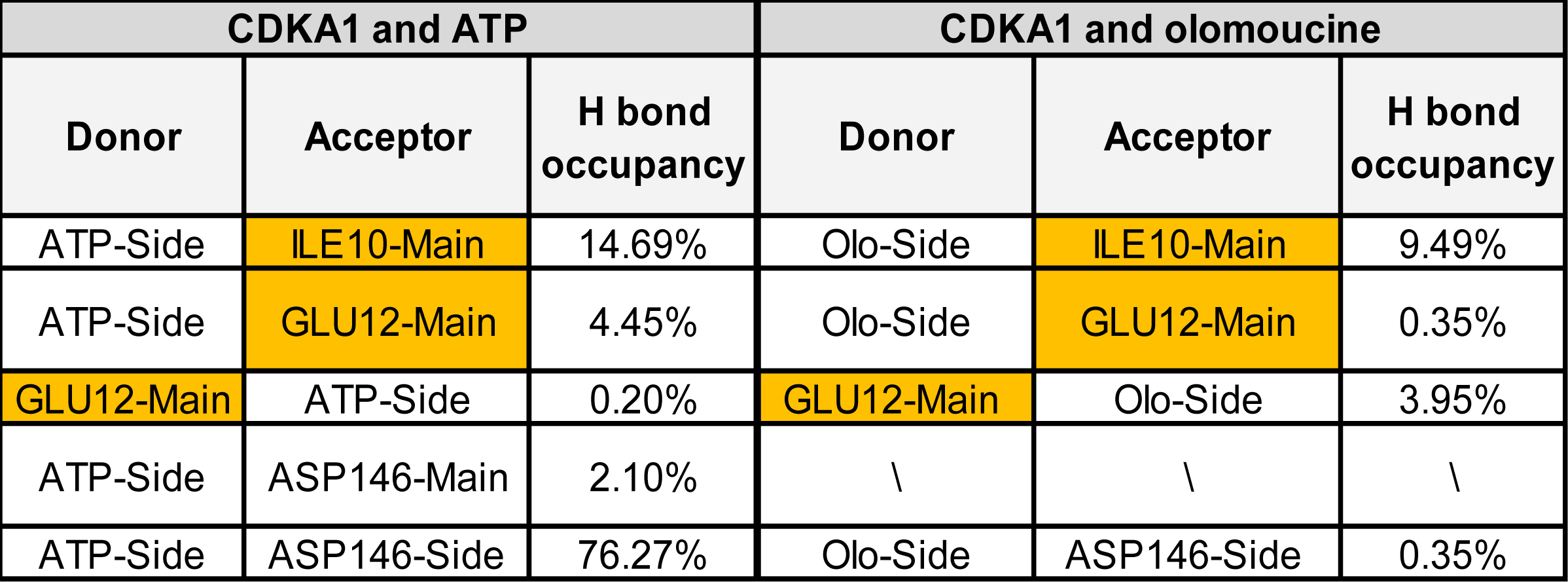

### Supplemental_figure12.tif

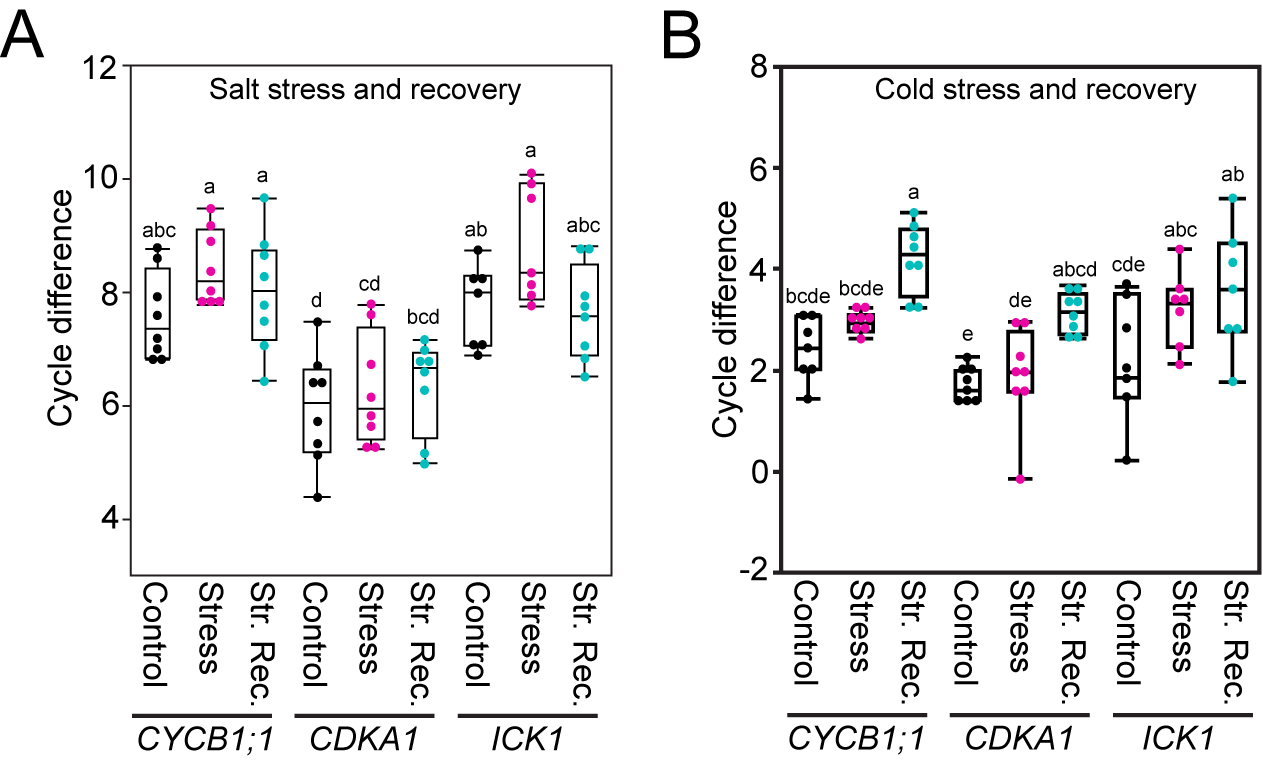
